# Partitioning heritability by functional category using GWAS summary statistics

**DOI:** 10.1101/014241

**Authors:** Hilary K. Finucane, Brendan Bulik-Sullivan, Alexander Gusev, Gosia Trynka, Yakir Reshef, Po-Ru Loh, Verneri Anttilla, Han Xu, Chongzhi Zang, Kyle Farh, Stephan Ripke, Felix Day, ReproGen Consortium, Schizophrenia Working Group of the Psychiatric Genetics Consortium, The RACI Consortium, Shaun Purcell, Eli Stahl, Sara Lindstrom, John R.B. Perry, Yukinori Okada, Soumya Raychaudhuri, Mark Daly, Nick Patterson, Benjamin M. Neale, Alkes L. Price

## Abstract

Recent work has demonstrated that some functional categories of the genome contribute disproportionately to the heritability of complex diseases. Here, we analyze a broad set of functional elements, including cell-type-specific elements, to estimate their polygenic contributions to heritability in genome-wide association studies (GWAS) of 17 complex diseases and traits spanning a total of 1.3 million phenotype measurements. To enable this analysis, we introduce a new method for partitioning heritability from GWAS summary statistics while controlling for linked markers. This new method is computationally tractable at very large sample sizes, and leverages genome-wide information. Our results include a large enrichment of heritability in conserved regions across many traits; a very large immunological disease-specific enrichment of heritability in FANTOM5 enhancers; and many cell-type-specific enrichments including significant enrichment of central nervous system cell types in body mass index, age at menarche, educational attainment, and smoking behavior. These results demonstrate that GWAS can aid in understanding the biological basis of disease and provide direction for functional follow-up.

## Introduction

In GWAS of complex traits, much of the heritability lies in single-nucleotide polymorphisms (SNPs) that do not reach genome-wide significance at current sample sizes.^1, 2^ However, many current approaches that leverage functional information^3, 4^ and GWAS data to inform disease biology use only SNPs in genome-wide significant loci,^5-7^ assume only one causal SNP per locus,^8^ or do not account for LD.^9^ We can improve power by estimating the proportion of SNP heritability^1^ attributable to various functional categories, using information from all SNPs and explicitly modeling LD.

Previous work on partitioning SNP heritability has used restricted maximum likelihood (REML) as implemented in GCTA.^10-13^ REML requires individual genotypes, but many of the largest GWAS analyses are conducted through meta-analysis of study-specific results, and so typically only summary statistics, not individual genotypes, are available for these studies. Even when individual genotypes are available, using REML to analyze multiple functional categories becomes computationally intractable at sample sizes in the tens of thousands. Here, we introduce a method for partitioning heritability, stratified LD score regression, that requires only GWAS summary statistics and LD information from an external reference panel that matches the population studied in the GWAS.

We apply our novel approach to 17 complex diseases and traits spanning 1,263,072 phenotype measurements. We first analyze non-cell-type-specific annotations and identify heritability enrichment in many of these functional annotations, including a large enrichment in conserved regions across many traits and a very large immunological disease-specific enrichment in FANTOM5 enhancers. We then analyze cell-type-specific annotations and identify many cell-type-specific heritability enrichments, including enrichment of central nervous system (CNS) cell types in body mass index, age at menarche, educational attainment, and smoking behavior.

## Results

### Overview of methods

Our method for partitioning heritability from summary statistics, called stratified LD score regression, relies on the fact that the *χ*^2^ association statistic for a given SNP includes the effects of all SNPs that it tags.^14, 15^ Thus, for a polygenic trait, SNPs with high linkage disequilibrium (LD) will have higher *χ*^2^ statistics on average than SNPs with low LD.^15^ This might be driven either by the higher likelihood of these SNPs to tag an individual large effect, or their ability to tag multiple weak effects. If we partition SNPs into functional categories with different contributions to heritability, then LD to a category that is enriched for heritability will increase the *χ*^2^ statistic of a SNP more than LD to a category that does not contribute to heritability. Thus, our method determines that a category of SNPs is enriched for heritability if SNPs with high LD to that category have higher *χ*^2^ statistics than SNPs with low LD to that category.

More precisely, under a polygenic model,^1^ the expected *χ*^2^ statistic of SNP *j* is

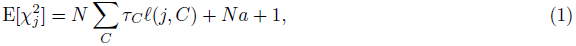

where *N* is sample size, *C* indexes disjoint categories, *ℓ*(*j*, *C*) is the LD Score of SNP *j* with respect to category *C* (defined as *ℓ*(*j*,*C*): = Σ_*kϵC*_*r*^*2*^(*j*,*k*)), *a* is a term that measures the contribution of confounding biases,^15^ and *τ*_C_ is the per-SNP heritability in category C (Methods). Equation (1) allows us to estimate *τ*_C_ via a (computationally simple) multiple regression of 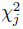against *ℓ*(*j*, *C*). The method easily generalizes to overlapping categories and case-control studies^16^ (Methods). We define the enrichment of a category to be the proportion of SNP heritability explained divided by the proportion of SNPs. We estimate standard errors with a block jackknife,^15^ and use these standard errors to calculate *z*-scores, *P*-values, and FDRs (Methods). We have released open-source software implementing the method (Web Resources).

To apply LD score regression (or REML) we must first specify which categories we include in our model. We created a “full baseline model” from 24 main annotations that are not specific to any cell type (Table S1; see Methods and Web Resources). Below, we show that including many categories in our model leads to more accurate estimates of enrichment. The 24 main annotations include: coding, UTR, promoter, and intron;^17^ histone marks H3K4me1, H3K4me3, H3K9ac^5^ and two versions of H3K27ac;^18, 19^ open chromatin reflected by DNase I hypersensitivity Site (DHS) regions;^5^ combined chromHMM/Segway predictions,^20^ which are a computational combination of many ENCODE annotations into a single partition of the genome into seven underlying “chromatin states,”; regions that are conserved in mammals;^21^ super-enhancers, which are large clusters of highly active enhancers;^19^ and active enhancers from the FANTOM5 panel of samples, which we call FANTOM5 enhancers.^22^ For the histone marks and other annotations that differ among cell types, we combined the different cell types into a single annotation for the baseline model by taking a union. To prevent our estimates from being biased upwards by enrichment in nearby regions,^13^ we also included 500bp windows around each functional category as separate functional categories in the baseline model, as well as 100bp windows around ChIP-seq peaks when appropriate (see Methods). This yielded a total of 53 (overlapping) functional categories in the baseline model, including a category containing all SNPs.

To prevent our estimates from being biased upwards by enrichment in nearby regions,^13^ we also included 500bp windows around each functional category as separate categories in the full baseline model, as well as 100bp windows around ChIP-seq peaks when appropriate (Methods). The 24 main annotations plus additional windows and a category containing all SNPs yielded a total of 53 (overlapping) functional categories in the full baseline model.

### Simulation results

In order to assess the robustness of the method, we performed a variety of simulations. We used chromosome 1 genotypes of controls from the Wellcome Trust Case Control Consortium,^23^ imputed using a 1000 Genomes reference panel.^24^ After quality control, we had 2,680 individuals and 360,106 SNPs (Methods). We simulated quantitative phenotypes with a heritability of 0.5 using an additive model. For each simulation, SNP effect sizes were drawn from a normal distribution with mean zero and variance (i.e., average per-SNP heritability) determined by functional categories. We estimated the enrichment of the DHS category, i.e., (%*h*^2^)/(%SNPs), using three methods:(1) REML with two categories (DHS/Other), (2) LD score regression with two categories (DHS/Other), and (3) LD score regression with the full baseline model (53 categories, described above). Since REML with 53 categories did not converge at this sample size and would be computationally intractable at sample sizes in the tens of thousands, we did not include it in our comparison; an advantage of LD score regression is that it is possible to include a large number of categories in the underlying model. We report means and standard errors of the mean over 100 independent simulations.

We first performed three sets of simulations where the causal pattern of enrichment was well modeled by the two-category (DHS/Other) model; for these, all three methods performed well, although LD score regression with the full baseline model had larger standard errors around the mean (Figure 1a). For example, the standard errors around the mean in simulations with no DHS enrichment were 0.08 for REML, 0.11 for two-category LD score regression and 0.19 for LD score regression with the full baseline model. For the first set of simulations, all SNPs were causal and SNP effect sizes were drawn i.i.d. from a normal distribution. For the second set of simulations, all SNPs were causal and SNP effect sizes were drawn independently from a normal distribution, but the variance of the normal distribution depended on whether the SNP was in a DHS region, and two variances (DHS and Other) were chosen so that the proportion of heritability of DHS would be 3x more than the proportion of SNPs. For the third set of simulations, only SNPs in DHS regions were causal, and effect sizes of DHS SNPs were drawn i.i.d. from a normal distribution.

**Figure 1:**
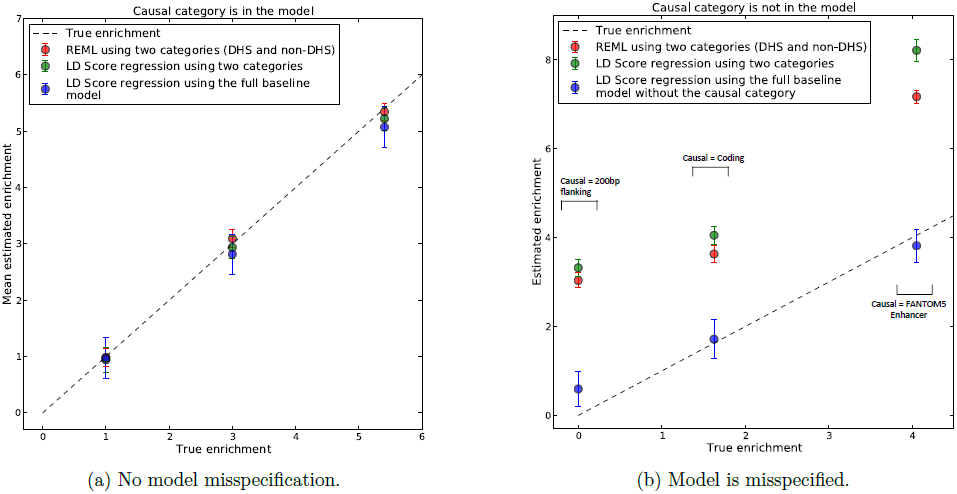
Simulation results. Enrichment is the proportion of heritability in DHS regions divided by the proportion of SNPs in DHS regions. Bars show 95% confidence intervals around the mean of 100 trials, (a) From left to right, the simulated genetic architectures are 1x DHS enrichment, 3x DHS enrichment, and 5.5x DHS enrichment (100% of heritability in DHS SNPs). (b) From left to right, the simulated genetic architectures are 200bp flanking regions causal, coding regions causal, and FANTOM5 Enhancer regions causal. For simulations with coding or FANTOM5 Enhancer as the causal category, we removed the causal category and the window around that category from the full baseline model in order to simulate enrichment in an unknown functional category.

Next, to explore the realistic scenario where the model used to estimate enrichment does not match the (unknown) causal model, we performed three sets of simulations where all causal SNPs were in a particular category, but the model used to estimate heritability did not include this causal category. The three sets of simulations were (1) all causal SNPs in coding regions, yielding 1.6x DHS enrichment due to coding/DHS overlap, (2) all causal SNPs in FANTOM5 enhancers, yielding 4.0x DHS enrichment due to FANTOM5 enhancer/DHS overlap, and (3) all causal SNPs in 200bp DHS flanking regions, yielding 0x DHS enrichment. For the coding and FANTOM5 enhancer causal simulations, we made the full baseline model into a misspecified model by removing the causal category (and windows around the causal category). Results from these simulations are displayed in Figure 1b).

The two-category estimators were not robust to model misspecification and consistently over-estimated DHS enrichment by a wide margin. LD score regression with the full baseline model gave more accurate mean estimates of enrichment. Specifically, for the simulations with coding and FANTOM5 Enhancers causal, LD score regression with the full baseline model gave unbiased mean enrichment estimates of 1.8x (s.e. 0.22) and 4.2x (s.e. 0.22), respectively, while the mean enrichment estimates of REML and two-category LD score regression were nearly double these. The full baseline model includes a 500bp window around DHS but not a 200bp window, and gave a mean estimated DHS enrichment of 0.65x (s.e. 0.22) when the 200bp flanking regions were causal, which is inflated relative to the true enrichment of 0x but much less inflated than > 3x mean enrichment estimates given by the two-category methods.

In summary, while these simulations include exaggerated patterns of enrichment (e.g., 100% of heritability in DHS flanking regions), the results highlight the possibility that two-category estimators of enrichment can yield incorrect conclusions. Although we cannot entirely rule out model misspecification as a source of bias for LD score regression with the full baseline model, we have shown here that it is robust to a wide variety of patterns of enrichment, because including many categories gives it the flexibility to adapt to the unknown causal model.

### Application to real data

We applied LD score regression with the full baseline model to 17 diseases and quantitative traits: height, BMI, age at menarche, LDL levels, HDL levels, triglyceride levels, coronary artery disease, type 2 diabetes, fasting glucose levels, schizophrenia, bipolar disorder, anorexia, educational attainment, smoking behavior, rheumatoid arthritis, Crohn’s disease, and ulcerative colitis (Table S2, Web resources). This includes all traits with publicly available summary statistics with sufficient sample size and SNP heritability, measured by the *z*-score of total SNP-heritability (Methods), spanning a total of of 1,263,072 unique phenotype measurements. We removed the MHC region from all analyses, due to its unusual LD and genetic architecture. Figure 2 shows results for the 24 main functional annotations, averaged across nine independent traits (Methods). Figure 3 shows trait-specific results for selected annotations and traits (Methods). Tables S3 and S4 show meta-analysis and trait-specific results for all traits and all 53 categories in the full baseline model.

We observed large and statistically significant enrichments for many functional categories. A few categories stood out in particular. First, regions conserved in mammals^21^ showed the largest enrichment of any category, with 2.6% of SNPs explaining an estimated 35% of SNP heritability on average across traits (*P* < 10^−15^ for enrichment). This is a significantly higher average enrichment than for coding regions, and provides evidence for the biological importance of conserved regions, even though the biochemical function of many conserved regions remains uncharacterized.^25^ Second, FANTOM5 Enhancers^22^ were extremely enriched in the three immunological diseases, with 0.4% of SNPs explaining an estimated 15% of SNP heritability on average across these three diseases (P < 10^−5^), but showed no evidence of enrichment for non-immunological traits (Figure 3). Third, repressed regions were depleted: 46% of SNPs explain only 29% of heritability on average (P < 0.006), consistent with the hypothesis that these are regions of low activity.^20^ We did not see a large enrichment of H3K27ac regions marked as super-enhancers over all H3K27ac regions; the estimates for enrichment were 1.8x (s.e. 0.2) and 1.6x (s.e. 0.1), respectively. This lack of enrichment supports the argument that super-enhancers may not play a much more important role in regulating transcription than regular enhancers.^26^ For many annotations, there was also enrichment in the 500bp flanking regions (Table S3). Analyses stratified by minor allele frequency produced broadly similar results for all of these enrichments (Table S5; see Methods).

**Figure 2:**
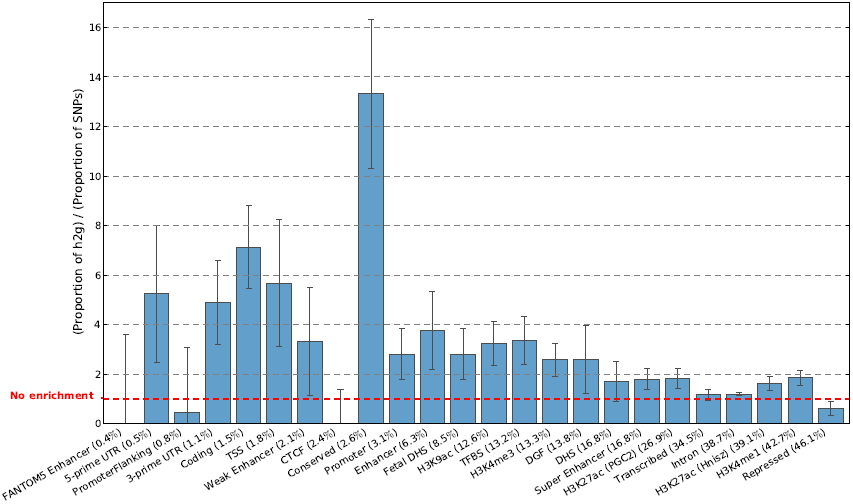
Enrichment estimates for the 24 main annotations, averaged over nine independent traits. Error bars represent 95% confidence intervals around the estimate.

**Figure 3:**
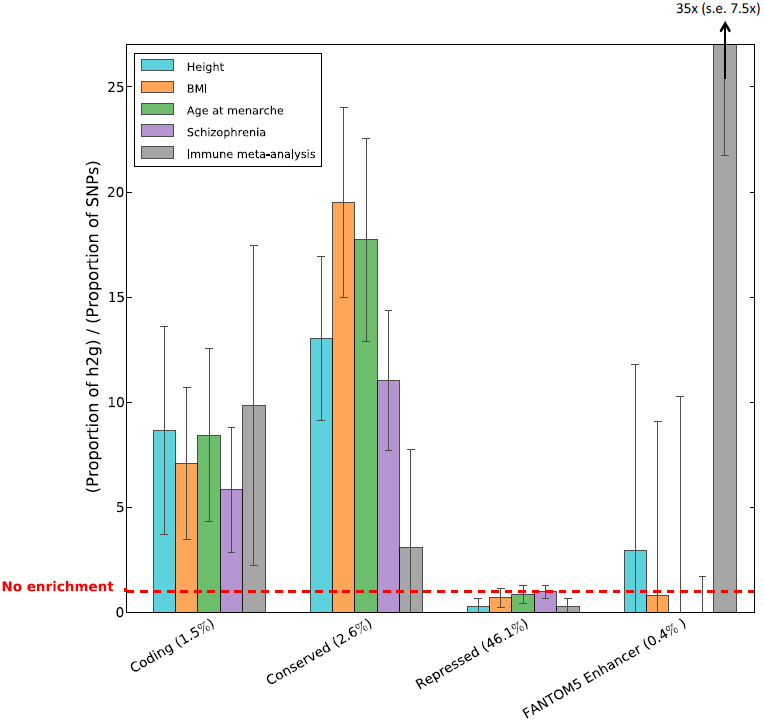
Enrichment estimates for selected annotations and traits. Error bars represent 95%confdence intervals around the estimate.

We performed two different cell-type-specific analyses: an analysis of 220 individual cell-type-specific annotations, and an analysis of 10 cell-type groups. The 220 individual cell-type-specific annotations are a combination of cell-type-specific annotations from four histone marks: 77 from H3K4me1,^5^ 81 from H3K4me3,^5^ 27 from H3K9ac,^5^ and 35 from H3K27ac^18^ (Table S6, Methods). When ranking these 220 cell-type-specific annotations, we wanted to control for overlap with the functional categories in the full baseline model, but not for overlap with the 219 other cell-type-specific annotations. Thus, we added the 220 cell-type-specific annotations individually, one at a time, to the full baseline model, and ranked these 220 annotations by the *P-value* for the coefficient corresponding to the annotation. This *P*-value tests whether the annotation contributes significantly to per-SNP heritability after controlling for the effects of the annotations in the full baseline model. We assessed statistical significance at the 0.05 level after Bonferroni correction for 220 × 17 = 3, 740 hypotheses tested. (This is conservative, since the 220 annotations are not independent.) We also report results with false discovery rate (FDR) < 0.05 (computed over 220 cell types for each trait). For 15 of the 17 traits, the top cell type passed an FDR threshold of 0.05. The top cell type for each trait is displayed in Table 1, with additional top cell types reported in Table S7.

**Table 1:** Enrichment of individual cell types. We report the cell type with the lowest *P*-value for each trait analyzed. * denotes FDR < 0.05. ** denotes significant at p < 0.05 after Bonferroni correction for multiple hypotheses. Sample sizes are inTable S2.

| Phenotype | Cell type | Tissue | Mark | $-\log_{10}(p)$ |
| --- | --- | --- | --- | --- |
| Height | Chondrogenic dif** | Bone | H3K27ac | 6.81 |
| BMI | Fetal brain* | Fetal brain | H3K4me3 | 4.48 |
| Age at menarche | Fetal brain** | Fetal brain | H3K4me3 | 12.25 |
| LDL | Liver (BI)* | Liver | H3K4me1 | 4.76 |
| HDL | Liver (BI)* | Liver | H3K4me1 | 4.51 |
| Triglycerides | Liver (BI)* | Liver | H3K4me1 | 3.99 |
| Coronary artery disease | Adipose nuclei* | Adipose | H3K4me1 | 4.21 |
| Type 2 Diabetes | Pancreatic islets | Pancreas | H3K4me3 | 2.87 |
| Fasting Glucose | Pancreatic islets* | Pancreas | H3K27ac | 3.93 |
| Schizophrenia | Fetal brain** | Fetal brain | H3K4me3 | 18.51 |
| Bipolar disorder | Mid frontal lobe* | Brain | H3K27ac | 4.42 |
| Anorexia | Angular gyrus | Brain | H3K9ac | 2.61 |
| Years of education | Angular gyrus** | Brain | H3K4me3 | 6.63 |
| Ever smoked | Inferior temporal lobe* | Brain | H3K4me3 | 3.21 |
| Rheumatoid arthritis | CD4+ CD25- IL17+ stim Th17** | Immune | H3K4me1 | 6.76 |
| Crohn's disease | CD4+ CD25- IL17+ stim Th17** | Immune | H3K4me1 | 7.59 |
| Ulcerative colitis | CD4+ CD25- IL17+ stim Th17** | Immune | H3K4me1 | 6.37 |

We combined information from related cell types by aggregating the 220 cell-type-specific annotations into 10 groups (Figure 4 legend and Table S6; see Methods). For each trait, we performed the same analysis on the 10 group-specific annotations as with the 220 cell-type-specific annotations. We assessed statistical significance at the 0.05 level after Bonferroni correction for 10 × 17 = 170 hypotheses tested, and we again also report results with false discovery rate (FDR) < 0.05 (now computed over all cell-type groups and traits). For 16 of the 17 traits (all traits except anorexia), the top cell type group passed an FDR threshold of 0.05. Results for the 11 traits with the most significant enrichments (after pruning closely related traits) are shown in Figure 4, with remaining traits in Figure S1.

**Figure 4:**
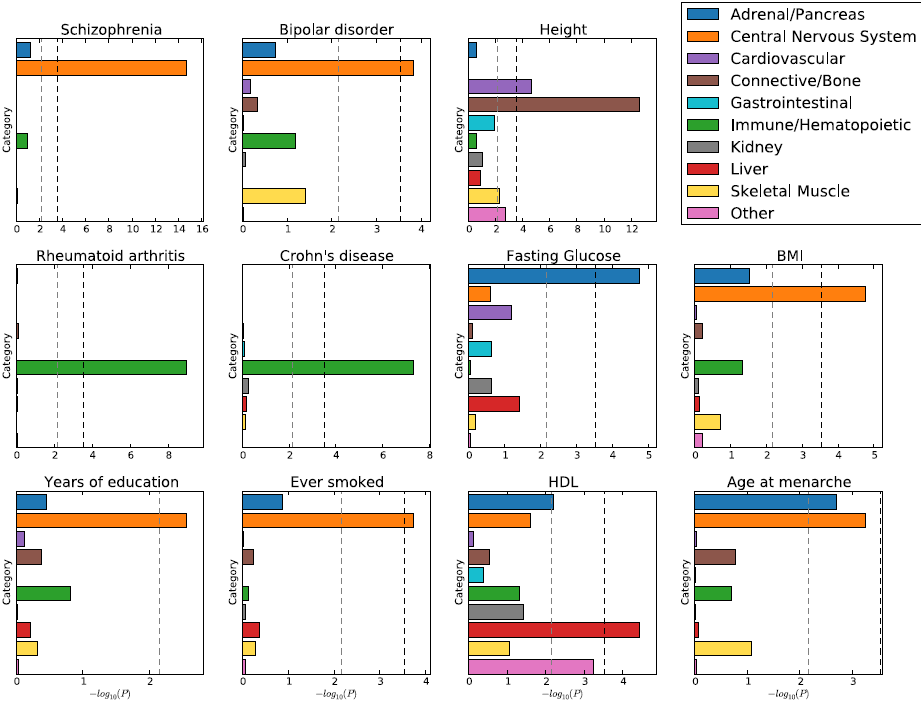
Enrichment of cell type groups. We report significance of enrichment for each of 10 cell-type groups, for each of 11 traits. The black dotted line at-log_10_(*P*) = 3:5 is the cutoff for Bonferroni significance. The grey dotted line at-log_10_(*P*) = 2:1 is the cutoff for FDR < 0.05. For HDL, three of the top individual cell types are adipose nuclei, which explains the enrichment of the “Other” category.

These two analyses are generally concordant, and show highly trait-specific patterns of cell-type enrichment. They also recapitulate several well-known findings. For example, the top cell type for each of the three lipid traits is liver (FDR < 0.05 for all three traits). For both type 2 diabetes and fasting glucose, the top cell type is pancreatic islets (FDR < 0.05 for fasting glucose but not type 2 diabetes). For the three psychiatric traits, the top cell type is a brain cell type (FDR < 0.05 for schizophrenia and bipolar disorder but not for anorexia) and the top cell-type group is CNS (significant after multiple testing for schizophrenia and bipolar disorder but not for anorexia).

There are also several new insights among these results. For example, the three immunological disorders show patterns of enrichment that reflect biological differences among the three disorders. Crohn’s disease has 40 cell types with FDR < 0.05, of which 39 are immune cell types and one (colonic mucosa) is a GI cell type. On the other hand, the 39 cell types with FDR < 0.05 for ulcerative colitis include nine GI cell types in addition to 30 immune cell types, whereas all 39 cell types with FDR < 0.05 for rheumatoid arthritis are immune cell types. The top cell type for all three traits is CD4+ CD25-IL17+ PMA Ionomycin simulated Th17 primary. Th17 cells are thought to act in opposition to Treg cells, which have been shown to suppress immune activity and whose malfunction has been associated with immunological disorders.^27^

We also identified several non-psychiatric phenotypes with enrichments in brain cell types. For both BMI and age at menarche, cell types in the central nervous system (CNS) ranked highest among individual cell types, and the top cell-type group was CNS, all with FDR < 0.05. These enrichments support previous human and animal studies that propose a strong neural basis for the regulation of energy homeostasis.^28^ For educational attainment, the top cell-type group is CNS (FDR < 0.05) and of the ten cell types that are significant after multiple testing, nine are CNS cell types. This is consistent with our understanding that the genetic component of educational attainment, which excludes environmental factors and population structure, is highly correlated with IQ.^29^ Finally, for smoking behavior, the CNS cell-type group is significant after multiple testing correction, and the top cell type is again a brain cell type, likely reflecting CNS involvement in nicotine processing.

## Discussion

We developed a new statistical method, stratified LD score regression, for identifying functional enrichment from GWAS data that uses information from all SNPs and explicitly models LD. We applied this method to GWAS data spanning 17 traits and 1.3 million phenotype measurements. Our method identified strong enrichment for conserved regions across all traits, and immunological disease-specific enrichment for FANTOM5 enhancers. Our cell-type-specific enrichment results confirmed previously known enrichments, such as liver enrichment for HDL levels and pancreatic islet enrichment for fasting glucose. In addition, we identified enrichments that would have been challenging to detect using existing methods, such as CNS enrichment for smoking behavior and educational attainment—traits with one and three genome-wide significant loci, respectively.^29, 30^ Stratified LD score regression represents a significant departure from previous methods that require raw genotypes,^10^ use only SNPs in genome-wide significant loci,^5-7^ assume only one causal SNP per locus,^8^ or do not account for LD^9^ (see Methods for a discussion of other methods). Our method is also computationally efficient, despite the 53 overlapping functional categories analyzed.

Although our polygenic approach has enabled a powerful analysis of genome-wide summary statistics, it has several limitations. First, the method requires a very large sample size and/or large SNP heritability, and the trait analyzed must be polygenic. Second, it requires an LD reference panel matched to the population studied; all results in this paper are from European datasets and use 1000G Europeans as a reference panel. Third, our method is currently not applicable to studies using custom genotyping arrays (e.g., Metabochip; see Methods). Fourth, our method is based on an additive model and does not consider the contribution of epistatic or other non-additive effects, nor does it model causal contributions of SNPs not in the reference panel; in particular, it is possible that patterns of enrichment may be different at extremely rare variants. Fifth, the method is limited by available functional data: if a trait is enriched in a cell type for which we have no data, we cannot detect the enrichment. Last, though we have shown our method to be robust in a wide range of scenarios, we cannot rule out model misspecification caused by enrichment in an unidentified functional category as a possible source of bias.

The polygenic approach described here is a powerful and efficient way to learn about functional enrichments from summary statistics, and it will become increasingly useful as functional data continues to grow and improve, and as GWAS studies of larger sample size are conducted.

## Web Resources

- ldsc software: https:github.com/bulik/ldsc
- DNaseI Digital Genomic Footprinting (DGF) annotations:^3, 13^ https://hgdownload.cse.ucsc.edu/goldenPath/hg19/encodeDCC/wgEncodeUwDgf
- Transcription factor binding sites:^3^ ^13^ https://hgdownload.cse.ucsc.edu/goldenPath/hg19/encodeDCC/wgEncodeAwgTfbsUniform/
- Segway-chromHMM combined enhancer annotations:^20^ ftp://ftp.ebi.ac.uk/pub/databases/ensembl/encode/integration_data_jan2011/byDataType/segmentations/jan2011
- Super-enhancers and H3K27ac: Available as a supplementary table to Hnisz et al 2014.^19^
- Conserved regions:^21, 31^ https://compbio.mit.edu/human-constraint/data/gff/">
- FANTOM5 Enhancers:^22^ https://enhancer.binf.ku.dk/presets/">
- Post-processed H3K4me1, H3K4me3, and H3K9ac:^5^ https://www.broadinstitute.org/mpg/goshifter/
- Height^32^ and BMI^33^ summary statistics: https://www.broadinstitute.org/collaboration/giant/index.php/GIANT_consortium_data_files
- Menarche summary statistics:^34^ https://www.reprogen.org
- LDL, HDL, and Triglycerides summary statistics:^35^ https://www.broadinstitute.org/mpg/pubs/lipids2010/
- Coronary artery disease summary statistics:^36^ https://www.cardiogramplusc4d.org
- Type 2 diabetes summary statistics:^37^ https://www.diagram-consortium.org
- Fasting glucose summary statistics:^38^ https://www.magicinvestigators.org/downloads/
- Schizophrenia,^18^ Bipolar Disorder,^39^ Anorexia,^40^ and Smoking behavior^30^ summary statistics: https://www.med.unc.edu/pgc/downloads
- Education attainment summary statistics:^29^ https://www.ssgac.org
- Rheumatoid arthritis summary statistics:^41^ https://plaza.umin.ac.jp/yokada/datasource/software.htm
- Crohn’s disease and ulcerative colitis summary statistics:^42^ https://www.ibdgenetics.org/downloads.html We used a newer version of these data with 1000 Genomes imputation.

## Acknowledgements

We thank Brad Bernstein, Mariel Finucane, Eran Hodis, Dylan Kotliar, X. Shirley Liu, Manolis Kellis, Michael O’Donovan, Bogdan Pasaniuc, Abhishek Sarkar, Patrick Sullivan, Bjarni Vilhjalmsson, and Adrian Veres for helpful discussions. This research was funded by NIH grants R01 MH101244, R03 CA173785, and 1U01HG0070033. H.K.F. was supported by the Fannie and John Hertz Foundation. S.R. is supported by funding from the Arthritis Foundation and by a Doris Duke Clinical Scientisit Development Award. This study made use of data generated by the Wellcome Trust Case Control Consortium (WTCCC) and the Wellcome Trust Sanger Institute. A full list of the investigators who contributed to the generation of the WTCCC data is available at www.wtccc.org.uk. Funding for the WTCCC project was provided by the Wellcome Trust under award 076113. The members of the Schizophrenia Working Group of the Psychiatric Genetics Consortium are listed in the Supplementary Information.

## Methods

### Stratified LD score regression

We begin with a derivation of Equation (1) for overlapping categories in a sample with no population structure or other confounding. The derivation of the intercept term in the presence of confounding is identical to the derivation in previous work.^15^

Let *y*_*i*_ be a quantitative phenotype in individual *i*, standardized to mean 0 and variance 1 in the population, and let *X*_*ij*_ be the genotype of individual *i* at the *j*-th SNP, standardized so that for each SNP *j*, *X*_*ij*_ has mean 0 and variance 1 in the population. We will assume a linear model:

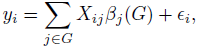

where *G* is some fixed set of SNPs, *β*_j_(*G*) is the effect size of SNP *j*, and *ϵ*_i_ is mean-0 noise.

We define *β*(*G*) = (*β*_1_(*G*),…, *β*_*M*_(*G*)) as the hypothetical result of multiple linear regression of *y* on *X* at infinite sample size. Thus, *β*(*G*) depends on the set *G*; for example, if *G* is the set of genotyped SNPs then *β*_*j*_(*G*) includes the causal effects of non-typed SNPs that are tagged by SNP *j*, whereas if *G* contains all SNPs, then *β*_*j*_(*G*) will reflect only the true effect at SNP *j*.

We will define the heritability of the set *G* of SNPs to be

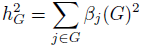

and the heritability of a category 𝓒 ⊂ *G* to be

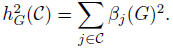

The definition of 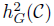
depends on both *G* and 𝓒; for example, if 𝓒 is the set of SNPs with minor allele frequency (MAF) greater than 5%, 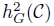 will be larger if G = 𝓒 than if *G* contains SNPs with lower MAF since in the first case 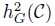 includes tagged effects of low-frequency SNPs, whereas in the second case the low-frequency effects are included in 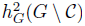.

Suppose that we have a sample of *N* individuals. Let *y* = (*y*_1_,…,*y*_*N*_), and let *X* be the *N* × *M* matrix of standardized genotypes, where *M* = |*G*|. (We will assume that our sample is large enough that normalizing each SNP within our sample is roughly equivalent to normalizing each SNP in the population.) Let *ϵ* = (*ϵ*_1_,…,*ϵ*_N_) be a vector of residuals. Then we can write

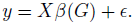

Let *β̂*_*j*_ be the estimate of the marginal effect of SNP *j* in our sample, given by

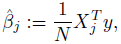

where *X*_*j*_ is the *j*-th column of *X*. Define *χ*^*2*^ statistics 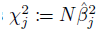.

Our goal here is to estimate 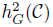, where *G* is the set of 1000G SNPs that have minor allele count greater than five in Europeans, and 𝓒 is a functional category, such as coding SNPs or SNPs that are in H3K4me3 regions in fetal brain tissue. From now on, we will omit the dependence on *G*, understanding *G* to be the set of 1000G SNPs with minor allele count greater than five in Europeans. We would like to estimate this quantity from summary statistics; i.e., the input to our method will be 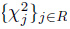, where *R* is the set of SNPs that were tested in a GWAS. We will also need LD information from a reference panel.

Substituting *y* = *Xβ* + *ϵ* into the the definition of *β̂*_j_, we get

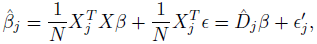

where *D̂*_*j*_ is the *j*-th row of the in-sample LD matrix *D̂* = *X*^*T*^*X/N* and 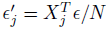. For a single entry *β*_*j*_, this means that

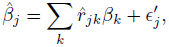

where *r̂*_*jk*_: = *D̂jk*) and 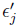 has mean 0 and variance 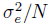.

To estimate 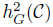, we will model *β* as a mean-0 random vector with independent entries. We will allow the variance of *β*_*j*_ to depend on certain functional properties of SNP *j*; for example, we will allow coding and non-coding SNPs to have different variances. More precisely, we will assume we have *C* functional categories 𝓒_1_,…, 𝓒_C_ ⊂ {1,…, *M*}. One of the categories must always contain all SNPs to allow for baseline heritability. We will model the variance of *β*_*j*_ as

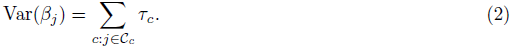

In the case that the 𝓒_c_ are disjoint, we will have *τ*_C_ = *h^2^*(*𝓒_c_*)/*M*(*𝓒_c_*), where *M*(*𝓒_c_*) is the number of SNPs in 𝓒_c_.

Consider the expectation of 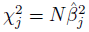.

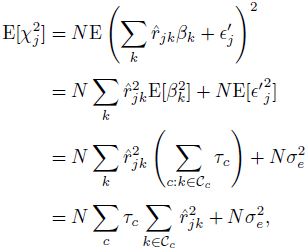

where the second equality follows because the random variables are all independent with mean 0.

Let *r*_*jk*_ denote the true correlation between SNPs *j* and *k* in the underlying population. In an unstructured sample 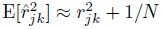.

We now have

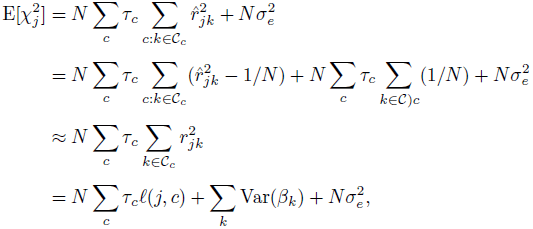

where 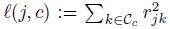. The variance of *y*_*j*_ is 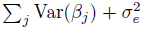. Since our phenotype has variance one, we can replace 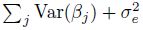 with 1, giving us our main equation:

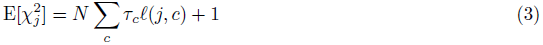

Our goal is to estimate 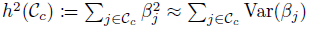, given a vector of *χ*^2^ statistics and LD information either from the sample or from a reference panel. To estimate Var(*β*_*j*_), we need to estimate each of the coefficients *τ*_*c*_, and then we apply Equation (2). To estimate *τ*_*c*_, we first compute *ℓ*(*j*, *c*), then we regress 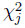on *ℓ*(*j*, *c*). This is called stratified LD score regression.

We estimate standard errors using a block jackknife over SNPs with 200 equally-sized blocks of adjacent SNPs,^15^ and use these standard errors to calculate *z*-scores, *P*-values, and FDRs. To minimize standard error, we remove outliers by excluding SNPs *j* with 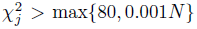, where *N* is the maximum sample size in the study, and we also weight the regression in a way that takes into account both over-counting and heteroskedasticity due to LD (see below). We remove the MHC region from all analyses.

#### Proportion of heritability and GC correction

As described above, stratified LD score regression is a method for estimating *h* ^2^ (𝓒) for a given category 𝓒. However, we are often more interested in estimating the proportion of heritability *h*^2^(𝓒)/*h*^2^. For this, we estimate *h*^2^(𝓒) and *h*^2^ separately and divide the estimates. For inference, we jackknife the proportion, rather than jackknifing the numerator and denominator separately.

Estimating the proportion of heritability is possible for summary statistics with GC correction, even though GC correction makes estimating the category specific heritability and total heritability impossible. This is because GC correction introduces a multiplicative error into estimates of both *h*^2^(𝓒) and *h*^2^,^15^ but the two multiplicative errors are equal, and cancel out in the ratio.

#### Choice of regression SNPs and reference SNPs

The derivation above does not incorporate imperfect imputation. Ideally, we would prune our *χ*^2^ statistics to a set of “regression SNPs” with imputation accuracy above 0.9, but since imputation accuracy is not always available, we instead use HapMap Project Phase 3 (HapMap3^43^) SNPs as a proxy for well-imputed SNPs. So for the purposes of this paper, our regression SNPs are always the HapMap3 SNPs.

However, we do not assume that only HapMap3 SNPs are causal. It is important that our model allow as many SNPs as possible to contribute causally, since if we use a model with, for example, only HapMap3 SNPs causal then we are assigning causality of any SNP that is tagged by HapMap3 but not included in HapMap3 to the HapMap3 SNPs that tag it. This is problematic specifically for functional partitioning because the functional categories containing the causal SNP may not be the same as the functional categories of the HapMap3 SNPs that tag it.

Recall that 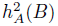 is the heritability of set *B* defined using a model that allows any SNP in set *A* to be causal. Another way to restate our above point is that we are interested in 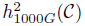 rather than 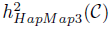 because a model that only allows HapMap3 SNPs to be causal is allowing non-HapMap3 heritability to be tagged by HapMap3 SNPs and therefore potentially assigning heritability to the wrong functional category. For this reason, our set of potentially causal SNPs—i.e., the set of SNPs in our reference panel—is the set of 1000G SNPs^24^ with minor allele count greater than five in Europeans.

However, there is a problem introduced by having many reference panel SNPs that are not tagged by regression SNPs: it leads us to extrapolate the enrichments at well-tagged SNPs to the rare SNPs on our reference panel that are not well-tagged. In other words, we estimate *τ*_*c*_ using summary statistics at HapMap3 SNPs, and then we multiply these coefficients by the number of SNPs in the relevant categories in 1000G, extrapolating the enrichments observed at HapMap3 SNPs to all 1000G SNPs.

To prevent ourselves from this extrapolation, we only estimate enrichments of categories containing common SNPs—i.e., SNPs with MAF > 0.05, all of which we assume to be well-tagged by HapMap3. Let 𝓖 denote the set of SNPs with *MAF* > 0.05. Then for any category 𝓒, we can estimate 𝓒 ∩ 𝓖 without potentially inaccurate extrapolation. For this reason, the proportions of heritability that we report throughout this manuscript are 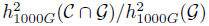.

#### Out-of-bounds estimates

Stratified LD score regression can give heritability estimates that are not between 0 and 1. When an unbiasedness is important—for example, when we are averaging estimates over several simulation replicates—we do not adjust these out-of-bounds estimates. However when mean squared error is more important than unbiasedness—for example, when reporting the results of a single analysis—we truncate these estimates to be between 0 and 1. To get a confidence interval around the truncated estimate, we intersect the original confidence interval with the interval [0, 1].

#### Custom genotyping arrays

LD score regression is not currently applicable to studies using a custom genotyping arrays. For these arrays, SNPs that are more likely to have a large effect size also have a larger sample size, and this dependency is not modeled in the above derivation.

#### Regression weights

To minimize standard error, we weight the regression in a way that takes into account both over-counting and heteroskedasticity due to LD. For over-counting, we compute LD Scores within HapMap3 SNPs; call these *ℓ*_*hm3*_(*j*). We also compute *ℓ*_1000*G*_(*j*, *c*) for all categories *c* in our model. The variance of 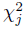 is proportional to (1 + *N* Σ_*c*_*τ*_*c*_*ℓ*_1000*G*_(*j*, *c*))^2^, but we do not have *τ*_*c*_. We use a rough approximation of *τ*_*c*_ obtained by taking the mean over regression SNPs of both sides of Equation (3) and assuming that all the *τ*_*c*_ are equal. This gives us *T̂* = (χ̄^2^ — 1) / (*N* . ℓ̄ (*j*)), where *χ*̄^2^ is the mean of *χ*^2^ and *Z*(*j*) is the mean of 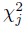 and *ℓ̄*(*j* is the mean of Σ_*c*_*ℓ*_1000*G*_(*j*, *c*), both taken over regression SNPs. We then weight SNP *j* by the inverse product of the over counting weights and heteroskedasticity weights:

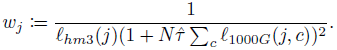

#### Baseline model

The 53 functional categories, derived from 24 main annotations, were obtained as follows:

- Coding, 3’-UTR, 5’-UTR promoter, and intron annotations from the RefSeq gene model were obtained from UCSC^17^ and post-processed by Gusev et al.^13^
- Digital genomic footprint and transcription factor binding site annotations were obtained from ENCODE^3^ and post-processed by Gusev et al.^13^
- The combined chromHMM/Segway annotations for six cell lines were obtained from Hoffman et al.^20^ CTCF, promoter flanking, transcribed, transcription start site, strong enhancer, and weak enhancers are a union the six cell lines; repressed is an intersection over the six cell lines.
- DNase I hypersensitive sites (DHSs) are a combination of ENCODE and Roadmap data, postprocessed by Trynka et al.^5^ We combined the cell-type-specific annotations into two annotations for inclusion in the full baseline model: a union of all cell types, and a union of only fetal cell types. DHS and fetal DHS.
- H3K4me1, H3K4me, and H3K9ac were all obtained from Roadmap and post-processed by Trynka et al.^5^ For each, we took a union over cell types for the full baseline model, and used the individual cell types for our cell-type-specific analysis.
- One version of H3K27ac was obtained from Roadmap, post-processed,^18^ and a second version of H3K27ac was obtained from the data of Hnisz et al.^19^ For each, we took a union over cell types for the full baseline model, and used the individual cell types for our cell-type-specific analysis.
- Super enhancers were also obtained from Hnisz et al,^19^ and comprise a subset of the H3K27ac annotation from that paper. We took a union over cell types for the full baseline model
- Regions conserved in mammals were obtained from Lindblad-Toh et al.,^21^ post-processed by Ward and Kellis.^31^
- FANTOM5 enhancers were obtained from Andersson et al.^22^
- For each of these 24 categories, we added a 500bp window around the category as an additional category to keep our heritability estimates from being inflated by heritability in flanking regions.^13^
- For each of DHS, H3K4me1, H3K4me3, and H3K9ac, we added a 100bp window around the ChIP-seq peak as an additional category.
- We added an additional category containing all SNPs.

#### WTCCC1 genotypes

We used chromosome 1 genotypes from the NBS and 1966BC control cohorts from the Wellcome Trust Case-Control Consortium.^23^ Imputation of the genotypes to integrated phase1 v3 1000 Genomes and QC were done as in Gusev et al.;^13^ we removed any SNPs that were below a MAF of 0.01, were above 0.002 missingness, or deviated from Hardy-Weinberg equilibrium at a *P* < 0.01.

#### Choice of phenotypes

We quantified the combination of sample size and heritability by the *z*-score of total SNP-heritability in the baseline analysis. We applied our method to all traits with available summary statistics, and removed all traits with a heritability *z*-score less than six. We then removed one of each pair of traits with a large genetic correlation (> 0.95): college attendance, which has a very high genetic correlation with years of education, and total cholesterol, which has a very high genetic correlation with LDL.^16^

#### Meta-analysis over traits

For our meta-analysis over traits, we identified pairs of traits with substantial sample overlap and trait correlation by using the intercept of LD score regression for genetic covariance. Specifically, we identified pairs of traits for which the intercept on N1N2 scale, which is an unbiased estimator of phenotypic correlation times sample overlap, was at least 10% of the sample size of either of the traits, and we excluded one of each such pair. The remaining set of traits was: Height, BMI, menarche, LDL levels, coronary artery disease, schizophrenia, educational attainment, smoking behavior, and rheumatoid arthritis. We then performed a random-effects meta-analysis of proportion of heritability over these phenotypes for each functional category. The results are in Figure 2 and Table S3.

#### Choice of traits to include in Figure 3

Height, BMI, age at menarche, and schizophrenia are the four traits with the highest combination of SNP heritability and sample size, which we quantify by the *z*-score of total heritability in the baseline analysis. We also included a meta-analysis of immunological diseases, since they have a different pattern of enrichment from other traits; for example FANTOM5 enhancers are very enriched for immunological diseases but not for other traits. This meta-analysis included only rheumatoid arthritis and Crohn’s disease; we excluded ulcerative colitis because that dataset shares controls with the Crohn’s disease dataset.

#### Robustness to derived allele frequency

To check that our results are not affected by a derived-allele-frequency-dependent genetic architecture, we repeated the meta-analysis over traits with a model that contained all of the categories of the full baseline model as well as seven derived allele frequency bins to the model as extra annotations: 0-0.1, 0.1-0.2, 0.2-0.3, 0.3-0.4, 0.4-0.6, 0.6-0.8, and 0.8-1. This allowed for effect size to depend on derived allele frequency, independently of functional annotation. These results are very similar to our results without derived allele frequency bins, and are displayed in Table S5.

#### Cell-type specific analysis

We used all available cell types from the four histone marks, excluding cell lines and cells labeled as cultured cells to limit ourselves to data with the clearest biological interpretation. The resulting cell-type-specific annotations are listed in Table S6. We then added each annotation individually to the baseline model and evaluated the significance of the coefficient *τ*_*c*_ of the cell-type-specific annofarh2014naturetation. Next, we combined the 220 cell-type-specific annotations into 10 cell-type groups and repeated the same analysis.

#### Discussion of other methods

There are no other methods designed to estimate genome-wide components of heritability from summary statistics. However, there are existing methods that identify enriched functional categories and cell types from summary statistics; here, we discuss a few of these methods.

A paper by Pickrell^8^ combines GWAS data with functional data to identify enriched and depleted functional categories, and leverages the resulting model to increase GWAS power. While the method is effective at increasing power and identifies many interesting enrichments, it is unclear whether the method is effective at ranking cell types; for example, fetal fibroblasts are the top cell type for Crohn’s disease. This could be because the model only allows for one causal SNP per locus, or could be a result of including all cell types simultaneously, thereby penalizing redundant cell types.

Kichaev et al.^7^ introduce a new method that leverages functional data for improved fine-mapping. Their method also outputs annotations associated with disease. While their method is effective in increasing fine-mapping resolution, it is again unclear whether the method is effective at ranking cell types; for example, the cell types they identify as contributing the most to HDL, LDL, and Triglycerides are muscle, kidney, and fetal small intestine, respectively, whereas the top cell types for those three phenotypes identified using our method are liver, liver, and liver. The lower power of this method in ranking cell types is likely because it considers only genome-wide significant loci. Similarly, a recent method from Farh et al.^6^ focuses on fine-mapping and considers only genome-wide significant loci, although their cell-type-specific results are more concordant with known biology.

Maurano et al. use enrichment of SNPs passing *P*-value thresholds of increasing stringency to identify important cell types. However, they are implicitly assuming that the functional annotation at a GWAS SNP matches the functional annotation at the causal SNP. While this could be true for functional annotations composed of very wide regions, it is not likely to be true for functional annotations composed of smaller regions, such as conserved regions. Moreover, their method does not account for total LD, and so could give biased results if used to compare functional annotations with different average amounts of total LD.

The method of Trynka et al.^5^ (also see a more recent manuscript^44^) is conservative in its identification of enrichment, comparing to a null obtained by local shifting rather than a genome-wide null, and leverages only genome-wide significant SNPs. As a result, they have very low power not only in traits such as bipolar disorder which have few genome-wide significant loci, but even for traits such as rheumatoid arthritis that have many significant loci. (Lowest *P*-value 10^−4^ much larger than the lowest *P*-value given by LD Score regression, 2 × 10^−7^.)

## Supplementary Information

### Members of the Schizophrenia Working Group of the Psychiatric Genetics Consortium

The members of the Schizophrenia Working Group of the Psychiatric Genetics Consortium are Stephan Ripke, Benjamin M. Neale, Aiden Corvin, James T.R. Walters, Kai-How Farh, Peter A. Holmans, Phil Lee, Brendan Bulik-Sullivan, David A. Collier, Hailiang Huang, Tune H. Pers, Ingrid Agartz, Esben Agerbo, Margot Albus, Madeline Alexander, Farooq Amin, Silviu A. Bacanu, Martin Begemann, Richard A. Belliveau, Jr., Judit Bene, Sarah E. Bergen, Elizabeth Bevilacqua, Tim B. Bigdeli, Donald W. Black, Anders D. Brglum, Richard Bruggeman, Nancy G. Buccola, Randy L. Buckner, William Byerley, Wiepke Cahn, Guiqing Cai, Dominique Campion, Rita M. Cantor, Vaughan J. Carr, Noa Carrera, Stanley V. Catts, Kimberly D. Chambert, Raymond C.K. Chan, Ronald Y.L. Chen, Eric Y.H. Chen, Wei Cheng, Eric F.C. Cheung, Siow Ann Chong, C. Robert Cloninger, David Cohen, Nadine Cohen, Paul Cormican, Nick Craddock, James J. Crowley, David Curtis, Michael Davidson, Kenneth L. Davis, Franziska Degenhardt, Jurgen Del Favero, Lynn E. DeLisi, Ditte Demontis, Dimitris Dikeos, Timothy Dinan, Srdjan Djurovic, Gary Donohoe, Elodie Drapeau, Jubao Duan, Frank Dudbridge, Naser Durmishi, Peter Eichhammer, Johan Eriksson, Valentina Escott-Price, Laurent Essioux, Ayman H. Fanous, Martilias S. Farrell, Josef Frank, Lude Franke, Robert Freedman, Nelson B. Freimer, Marion Friedl, Joseph I. Friedman, Menachem Fromer, Giulio Genovese, Lyudmila Georgieva, Elliot S. Gershon, Ina Giegling, Paola Giusti-Rodrguez, Stephanie Godard, Jacqueline I. Goldstein, Vera Golimbet, Srihari Gopal, Jacob Gratten, Jakob Grove, Lieuwe de Haan, Christian Hammer, Marian L. Hamshere, Mark Hansen, Thomas Hansen, Vahram Haroutunian, Annette M. Hartmann, Frans A. Henskens, Stefan Herms, Joel N. Hirschhorn, Per Hoffmann, Andrea Hofman, Mads V. Hollegaard, David M. Hougaard, Masashi Ikeda, Inge Joa, Antonio Julia, Rene S. Kahn, Luba Kalaydjieva, Sena Karachanak-Yankova, Juha Karjalainen, David Kavanagh, Matthew C. Keller, Brian J. Kelly, James L. Kennedy, Andrey Khrunin, Yunjung Kim, Janis Klovins, James A. Knowles, Bettina Konte, Vaidutis Kucinskas, Zita Ausrele Kucinskiene, Hana Kuzelova-Ptackova, Anna K. Kahler, Claudine Laurent, Jimmy Lee Chee Keong, S. Hong Lee, Sophie E. Legge, Bernard Lerer, Miaoxin Li, Tao Li, Kung-Yee Liang, Jeffrey Lieberman, Svetlana Limborska, Carmel M. Loughland, Jan Lubinski, Jouko Lnnqvist, Milan Macek, Jr., Patrik K.E. Magnusson, Brion S. Maher, Wolfgang Maier, Jacques Mallet, Sara Marsal, Manuel Mattheisen, Morten Mattingsdal, Robert W. McCarley, Colm McDonald, Andrew M. McIntosh, Sandra Meier, Carin J. Meijer, Bela Melegh, Ingrid Melle, Raquelle I. Mesholam-Gately, Andres Metspalu, Patricia T. Michie, Lili Milani, Vihra Milanova, Younes Mokrab, Derek W. Morris, Ole Mors, Preben B. Mortensen, Kieran C. Murphy, Robin M. Murray, Inez Myin-Germeys, Bertram Mller-Myhsok, Mari Nelis, Igor Nenadic, Deborah A. Nertney, Gerald Nestadt, Kristin K. Nicodemus, Liene Nikitina-Zake, Laura Nisenbaum, Annelie Nordin, Eadbhard O’Callaghan, Colm O’Dushlaine, F. Anthony O’Neill, Sang-Yun Oh, Ann Olincy, Line Olsen, Jim Van Os, Psychosis Endophenotypes International Consortium, Christos Pantelis, George N. Papadimitriou, Sergi Papiol, Elena Parkhomenko, Michele T. Pato, Tiina Paunio, Milica Pejovic-Milovancevic, Diana O. Perkins, Olli Pietilinen, Jonathan Pimm, Andrew J. Pocklington, John Powell, Alkes Price, Ann E. Pulver, Shaun M. Purcell, Digby Quested, Henrik B. Rasmussen, Abraham Reichenberg, Mark A. Reimers, Alexander L. Richards, Joshua L. Roffman, Panos Roussos, Douglas M. Ruderfer, Veikko Salomaa, Alan R. Sanders, Ulrich Schall, Christian R. Schubert, Thomas G. Schulze, Sibylle G. Schwab, Edward M. Scolnick, Rodney J. Scott, Larry J. Seidman, Jianxin Shi, Engilbert Sigurdsson, Teimuraz Silagadze, Jeremy M. Silverman, Kang Sim, Petr Slominsky, Jordan W. Smoller, Hon-Cheong So, Chris C.A. Spencer, Eli A. Stahl, Hreinn Stefansson, Stacy Steinberg, Elisabeth Stogmann, Richard E. Straub, Eric Strengman, Jana Strohmaier, T. Scott Stroup, Mythily Subramaniam, Jaana Suvisaari, Dragan M. Svrakic, Jin P. Szatkiewicz, Erik Sderman, Srinivas Thirumalai, Draga Toncheva, Paul A. Tooney, Sarah Tosato, Juha Veijola, John Waddington, Dermot Walsh, Dai Wang, Qiang Wang, Bradley T. Webb, Mark Weiser, Dieter B. Wildenauer, Nigel M. Williams, Stephanie Williams, Stephanie H. Witt, Aaron R. Wolen, Emily H.M. Wong, Brandon K. Wormley, Jing Qin Wu, Hualin Simon Xi, Clement C. Zai, Xuebin Zheng, Fritz Zimprich, Naomi R. Wray, Kari Stefansson, Peter M. Visscher, Wellcome Trust Case Control Consortium, Rolf Adolfsson, Ole A. Andreassen, Douglas H.R. Blackwood, Elvira Bramon, Joseph D. Buxbaum, Anders D. Brglum, Sven Cichon, Ariel Darvasi, Enrico Domenici, Hannelore Ehrenreich, Tonu Esko, Pablo V. Gejman, Michael Gill, Hugh Gurling, Christina M. Hultman, Nakao Iwata, Assen V. Jablensky, Erik G. Jons-son, Kenneth S. Kendler, George Kirov, Jo Knight, Todd Lencz, Douglas F. Levinson, Qingqin S. Li, Jianjun Liu, Anil K. Malhotra, Steven A. McCarroll, Andrew McQuillin, Jennifer L. Moran, Preben B. Mortensen, Bryan J. Mowry, Markus M. Nthen, Roel A. Ophoff, Michael J. Owen, Aarno Palotie, Carlos N. Pato, Tracey L. Petryshen, Danielle Posthuma, Marcella Rietschel, Brien P. Riley, Dan Rujescu, Pak C. Sham, Pamela Sklar, David St. Clair, Daniel R. Weinberger, Jens R. Wendland, Thomas Werge, Mark J. Daly, Patrick F. Sullivan, and Michael C. O’Donovan.

**Table S1:** Annotations used. For DHS, H3K4me1, H3K4me3, and H3K9ac, we include peaks and regions as two annotations. For the annotations from the Hoffman segmentation,^*20*^ we union over six cell lines for each category except Repressed, where we take an intersection instead. We also include a 500bp window around each annotation as a separate annotation in the model.

| Annotation | Prop. SNPs | Mean segment length (bp) |
| --- | --- | --- |
| Coding | 0.015 | 315 |
| Conserved | 0.026 | 34 |
| CTCF | 0.024 | 490 |
| DGF | 0.138 | 208 |
| DHS | 0.168 | 358 |
| FANTOM5 Enhancer | 0.004 | 289 |
| Enhancer | 0.063 | 678 |
| Fetal DHS | 0.085 | 339 |
| H3K27ac<sup>19</sup> | 0.391 | 12411 |
| H3K27ac<sup>18</sup> | 0.269 | 1051 |
| H3K4me1 | 0.427 | 1676 |
| H3K4me3 | 0.133 | 941 |
| H3K9ac | 0.126 | 964 |
| Intron | 0.387 | 6537 |
| PromoterFlanking | 0.008 | 266 |
| Promoter | 0.031 | 4192 |
| Repressed | 0.461 | 572 |
| Super Enhancer | 0.168 | 54744 |
| TFBS | 0.132 | 509 |
| Transcribed | 0.345 | 484 |
| TSS | 0.018 | 813 |
| 3-prime UTR | 0.011 | 844 |
| 5-prime UTR | 0.005 | 197 |
| Weak Enhancer | 0.021 | 249 |

**Table S2:** Phenotypes used in the main analyses. The total number of phenotype measurements is 1,263,072. There is sample overlap among the studies, so the number of unique individuals is lower.

| Phenotype | Reference/consortium | *N* |
| --- | --- | --- |
| Height | Lango Allen et al., 2010 Nature | 133,858 |
| BMI | Speliotes et al., 2010 Nat Genet | 123,912 |
| Age at menarche | Perry et al., 2014 Nature | 132,989 |
| LDL | Teslovich et al., 2010 Nature | 95,454 |
| HDL | Teslovich et al., 2010 Nature | 99,900 |
| Triglycerides | Teslovich et al., 2010 Nature | 96,598 |
| Coronary Artery Disease | Schunkert et al., Nat Genet 2011 | 86,995 |
| Type-2 Diabetes | Morris et al., 2012 Nat Genet | 69,033 |
| Fasting glucose | Manning et. al., Nat Genet, 2012 | 58,074 |
| Schizophrenia | SCZ Working Group of the PGC, 2014 Nature | 70,100 |
| Bipolar Disorder | Bip Working Group of the PGC, 2011 Nat Genet | 16,731 |
| Anorexia | Boraska et al., 2014 Mol Psych | 17,767 |
| Educational attainment | Rietveld et al., Science 2013 | 101,069 |
| Ever smoked | TAG Consortium, 2010 Nat Genet | 74,035 |
| Rheumatoid Arthritis | Okada et al., 2014 Nature | 38,242 |
| Crohn's Disease | Jostins et al., 2012 Nature | 20,883 |
| Ulcerative Colitis | Jostins et al., 2012 Nature | 27,432 |

**Table S3:** Proportion of SNP-heritability and enrichment for different functional categories. We display results meta-analyzed across nine traits for each of the 53 annotations, including two distinct H3K27ac annotations (Methods). Although true SNP-heritability is non-negative, we report here unbiased estimates, we can be negative (Methods).

| Annotation | Prop. SNPs | Prop. $h^2$ | Enrichment |
| --- | --- | --- | --- |
| Coding | 0.015 | 0.104 (0.012) | 7.122 (0.840) |
| Coding + 500bp | 0.065 | 0.190 (0.030) | 2.938 (0.467) |
| Conserved | 0.026 | 0.347 (0.039) | 13.322 (1.500) |
| Conserved + 500bp | 0.333 | 0.654 (0.026) | 1.966 (0.078) |
| CTCF | 0.024 | -0.004 (0.019) | -0.176 (0.789) |
| CTCF + 500bp | 0.071 | 0.059 (0.019) | 0.825 (0.272) |
| DGF | 0.138 | 0.358 (0.094) | 2.604 (0.686) |
| DGF + 500bp | 0.542 | 0.761 (0.069) | 1.405 (0.128) |
| DHS peaks | 0.112 | 0.224 (0.063) | 2.005 (0.566) |
| DHS | 0.168 | 0.285 (0.069) | 1.699 (0.410) |
| DHS + 500bp | 0.499 | 0.788 (0.041) | 1.580 (0.082) |
| FANTOM5 Enhancer | 0.004 | -0.003 (0.009) | -0.727 (2.160) |
| FANTOM5 Enhancer + 500bp | 0.019 | 0.017 (0.017) | 0.878 (0.896) |
| Enhancer | 0.063 | 0.238 (0.050) | 3.764 (0.784) |
| Enhancer + 500bp | 0.154 | 0.359 (0.042) | 2.334 (0.272) |
| Fetal DHS | 0.085 | 0.238 (0.043) | 2.805 (0.512) |
| Fetal DHS + 500bp | 0.285 | 0.591 (0.058) | 2.073 (0.204) |
| H3K27ac<sup>19</sup> | 0.391 | 0.631 (0.054) | 1.612 (0.138) |
| H3K27ac<sup>19</sup> + 500bp | 0.423 | 0.664 (0.059) | 1.572 (0.140) |
| H3K27ac<sup>18</sup> | 0.269 | 0.490 (0.054) | 1.818 (0.200) |
| H3K27ac<sup>18</sup> + 500bp | 0.336 | 0.611 (0.040) | 1.817 (0.118) |
| H3K4me1 peaks | 0.171 | 0.447 (0.040) | 2.611 (0.235) |
| H3K4me1 | 0.427 | 0.792 (0.065) | 1.857 (0.152) |
| H3K4me1 + 500bp | 0.609 | 0.910 (0.039) | 1.494 (0.064) |
| H3K4me3 peaks | 0.042 | 0.158 (0.025) | 3.780 (0.599) |
| H3K4me3 | 0.133 | 0.344 (0.045) | 2.582 (0.336) |
| H3K4me3 + 500bp | 0.255 | 0.487 (0.056) | 1.905 (0.220) |
| H3K9ac peaks | 0.039 | 0.248 (0.024) | 6.399 (0.618) |
| H3K9ac | 0.126 | 0.408 (0.056) | 3.233 (0.441) |
| H3K9ac + 500bp | 0.231 | 0.504 (0.040) | 2.184 (0.172) |
| Intron | 0.387 | 0.462 (0.014) | 1.192 (0.035) |
| Intron + 500bp | 0.397 | 0.521 (0.015) | 1.313 (0.037) |
| PromoterFlanking | 0.008 | 0.004 (0.011) | 0.448 (1.314) |
| PromoterFlanking + 500bp | 0.033 | 0.081 (0.018) | 2.433 (0.531) |
| Promoter | 0.031 | 0.087 (0.016) | 2.806 (0.512) |
| Promoter + 500bp | 0.039 | 0.080 (0.016) | 2.063 (0.424) |
| Repressed | 0.461 | 0.286 (0.063) | 0.619 (0.137) |
| Repressed + 500bp | 0.719 | 0.446 (0.049) | 0.620 (0.068) |
| Super Enhancer | 0.168 | 0.304 (0.035) | 1.804 (0.210) |
| Super Enhancer + 500bp | 0.172 | 0.319 (0.037) | 1.857 (0.216) |
| TFBS | 0.132 | 0.445 (0.063) | 3.357 (0.479) |
| TFBS + 500bp | 0.343 | 0.503 (0.052) | 1.464 (0.152) |
| Transcribed | 0.345 | 0.407 (0.038) | 1.179 (0.111) |
| Transcribed + 500bp | 0.763 | 0.721 (0.028) | 0.945 (0.036) |
| TSS | 0.018 | 0.104 (0.023) | 5.682 (1.278) |
| TSS + 500bp | 0.035 | 0.172 (0.029) | 4.940 (0.827) |
| 3-prime UTR | 0.011 | 0.054 (0.009) | 4.892 (0.856) |
| 3-prime UTR + 500bp | 0.027 | 0.074 (0.011) | 2.730 (0.412) |
| 5-prime UTR | 0.005 | 0.028 (0.008) | 5.243 (1.385) |
| 5-prime UTR + 500bp | 0.028 | 0.065 (0.010) | 2.340 (0.374) |
| Weak Enhancer | 0.021 | 0.070 (0.023) | 3.323 (1.090) |
| Weak Enhancer + 500bp | 0.089 | 0.199 (0.030) | 2.238 (0.335) |

**(S4.A).** Proportion of heritability and enrichment for different functional categories for Height.

| Annotation | Prop. SNPs | Prop. $h^2$ | Enrichment |
| --- | --- | --- | --- |
| Coding | 0.015 | 0.127 (0.036) | 8.661 (2.475) |
| Coding + 500bp | 0.065 | 0.192 (0.034) | 2.979 (0.521) |
| Conserved | 0.026 | 0.340 (0.051) | 13.029 (1.950) |
| Conserved + 500bp | 0.333 | 0.636 (0.053) | 1.913 (0.159) |
| CTCF | 0.024 | 0.050 (0.045) | 2.118 (1.909) |
| CTCF + 500bp | 0.071 | 0.089 (0.048) | 1.249 (0.669) |
| DGF | 0.138 | 0.585 (0.085) | 4.253 (0.621) |
| DGF + 500bp | 0.542 | 0.983 (0.066) | 1.815 (0.122) |
| DHS peaks | 0.112 | 0.429 (0.107) | 3.837 (0.958) |
| DHS | 0.168 | 0.512 (0.120) | 3.050 (0.717) |
| DHS + 500bp | 0.499 | 0.787 (0.062) | 1.579 (0.124) |
| FANTOM5 Enhancer | 0.004 | 0.013 (0.019) | 2.930 (4.429) |
| FANTOM5 Enhancer + 500bp | 0.019 | 0.014 (0.023) | 0.711 (1.224) |
| Enhancer | 0.063 | 0.184 (0.064) | 2.906 (1.018) |
| Enhancer + 500bp | 0.154 | 0.396 (0.054) | 2.573 (0.353) |
| Fetal DHS | 0.085 | 0.281 (0.089) | 3.311 (1.050) |
| Fetal DHS + 500bp | 0.285 | 0.504 (0.080) | 1.770 (0.280) |
| H3K27ac<sup>19</sup> | 0.391 | 0.782 (0.035) | 1.999 (0.089) |
| H3K27ac<sup>19</sup> + 500bp | 0.423 | 0.780 (0.043) | 1.846 (0.101) |
| H3K27ac<sup>18</sup> | 0.269 | 0.533 (0.065) | 1.978 (0.243) |
| H3K27ac<sup>18</sup> + 500bp | 0.336 | 0.674 (0.061) | 2.005 (0.181) |
| H3K4me1 peaks | 0.171 | 0.422 (0.091) | 2.465 (0.530) |
| H3K4me1 | 0.427 | 0.965 (0.074) | 2.261 (0.173) |
| H3K4me1 + 500bp | 0.609 | 0.946 (0.036) | 1.553 (0.060) |
| H3K4me3 peaks | 0.042 | 0.075 (0.053) | 1.791 (1.263) |
| H3K4me3 | 0.133 | 0.429 (0.071) | 3.215 (0.535) |
| H3K4me3 + 500bp | 0.255 | 0.584 (0.054) | 2.286 (0.211) |
| H3K9ac peaks | 0.039 | 0.266 (0.060) | 6.851 (1.539) |
| H3K9ac | 0.126 | 0.558 (0.066) | 4.428 (0.520) |
| H3K9ac + 500bp | 0.231 | 0.602 (0.055) | 2.612 (0.237) |
| Intron | 0.387 | 0.448 (0.031) | 1.156 (0.081) |
| Intron + 500bp | 0.397 | 0.532 (0.025) | 1.340 (0.064) |
| PromoterFlanking | 0.008 | -0.046 (0.025) | -5.400 (3.021) |
| PromoterFlanking + 500bp | 0.033 | 0.116 (0.034) | 3.453 (1.029) |
| Promoter | 0.031 | 0.089 (0.033) | 2.848 (1.057) |
| Promoter + 500bp | 0.039 | 0.101 (0.020) | 2.603 (0.528) |
| Repressed | 0.461 | 0.115 (0.093) | 0.248 (0.203) |
| Repressed + 500bp | 0.719 | 0.369 (0.039) | 0.513 (0.054) |
| Super Enhancer | 0.168 | 0.384 (0.026) | 2.280 (0.155) |
| Super Enhancer + 500bp | 0.172 | 0.398 (0.023) | 2.317 (0.137) |
| TFBS | 0.132 | 0.455 (0.083) | 3.438 (0.628) |
| TFBS + 500bp | 0.343 | 0.682 (0.065) | 1.984 (0.190) |
| Transcribed | 0.345 | 0.572 (0.094) | 1.657 (0.272) |
| Transcribed + 500bp | 0.763 | 0.652 (0.052) | 0.854 (0.068) |
| TSS | 0.018 | 0.107 (0.038) | 5.853 (2.103) |
| TSS + 500bp | 0.035 | 0.174 (0.032) | 4.991 (0.905) |
| 3-prime UTR | 0.011 | 0.102 (0.032) | 9.240 (2.857) |
| 3-prime UTR + 500bp | 0.027 | 0.118 (0.032) | 4.380 (1.176) |
| 5-prime UTR | 0.005 | 0.045 (0.022) | 8.310 (4.056) |
| 5-prime UTR + 500bp | 0.028 | 0.069 (0.023) | 2.470 (0.840) |
| Weak Enhancer | 0.021 | 0.018 (0.043) | 0.853 (2.025) |
| Weak Enhancer + 500bp | 0.089 | 0.109 (0.048) | 1.229 (0.539) |

**(S4.B).** Proportion of heritability and enrichment for different functional categories for BMI.

(S4.A) Proportion of heritability and enrichment for different functional categories for Height.
| Annotation | Prop. SNPs | Prop. $h^2$ | Enrichment |
| --- | --- | --- | --- |
| Coding | 0.015 | 0.104 (0.027) | 7.071 (1.813) |
| Coding + 500bp | 0.065 | 0.129 (0.032) | 1.992 (0.500) |
| Conserved | 0.026 | 0.508 (0.059) | 19.504 (2.270) |
| Conserved + 500bp | 0.333 | 0.545 (0.068) | 1.640 (0.205) |
| CTCF | 0.024 | -0.027 (0.049) | -1.135 (2.051) |
| CTCF + 500bp | 0.071 | 0.025 (0.045) | 0.358 (0.640) |
| DGF | 0.138 | 0.034 (0.124) | 0.249 (0.899) |
| DGF + 500bp | 0.542 | 0.674 (0.068) | 1.244 (0.125) |
| DHS peaks | 0.112 | 0.125 (0.100) | 1.123 (0.894) |
| DHS | 0.168 | 0.093 (0.123) | 0.554 (0.733) |
| DHS + 500bp | 0.499 | 0.890 (0.123) | 1.785 (0.246) |
| FANTOM5 Enhancer | 0.004 | 0.003 (0.018) | 0.792 (4.139) |
| FANTOM5 Enhancer + 500bp | 0.019 | -0.004 (0.026) | -0.218 (1.364) |
| Enhancer | 0.063 | 0.259 (0.052) | 4.083 (0.829) |
| Enhancer + 500bp | 0.154 | 0.305 (0.052) | 1.980 (0.337) |
| Fetal DHS | 0.085 | 0.112 (0.089) | 1.320 (1.045) |
| Fetal DHS + 500bp | 0.285 | 0.572 (0.081) | 2.008 (0.284) |
| H3K27ac<sup>19</sup> | 0.391 | 0.514 (0.037) | 1.315 (0.096) |
| H3K27ac<sup>19</sup> + 500bp | 0.423 | 0.564 (0.039) | 1.334 (0.093) |
| H3K27ac<sup>18</sup> | 0.269 | 0.447 (0.059) | 1.660 (0.221) |
| H3K27ac<sup>18</sup> + 500bp | 0.336 | 0.557 (0.067) | 1.657 (0.199) |
| H3K4me1 peaks | 0.171 | 0.394 (0.102) | 2.298 (0.595) |
| H3K4me1 | 0.427 | 0.668 (0.092) | 1.565 (0.216) |
| H3K4me1 + 500bp | 0.609 | 0.880 (0.052) | 1.445 (0.086) |
| H3K4me3 peaks | 0.042 | 0.150 (0.054) | 3.583 (1.295) |
| H3K4me3 | 0.133 | 0.219 (0.057) | 1.646 (0.429) |
| H3K4me3 + 500bp | 0.255 | 0.405 (0.054) | 1.585 (0.212) |
| H3K9ac peaks | 0.039 | 0.279 (0.056) | 7.206 (1.452) |
| H3K9ac | 0.126 | 0.214 (0.066) | 1.699 (0.520) |
| H3K9ac + 500bp | 0.231 | 0.450 (0.051) | 1.952 (0.222) |
| Intron | 0.387 | 0.424 (0.032) | 1.094 (0.083) |
| Intron + 500bp | 0.397 | 0.490 (0.030) | 1.233 (0.076) |
| PromoterFlanking | 0.008 | 0.004 (0.023) | 0.477 (2.702) |
| PromoterFlanking + 500bp | 0.033 | 0.042 (0.035) | 1.262 (1.035) |
| Promoter | 0.031 | 0.080 (0.031) | 2.559 (0.989) |
| Promoter + 500bp | 0.039 | 0.060 (0.024) | 1.565 (0.615) |
| Repressed | 0.461 | 0.321 (0.105) | 0.696 (0.227) |
| Repressed + 500bp | 0.719 | 0.619 (0.032) | 0.860 (0.045) |
| Super Enhancer | 0.168 | 0.217 (0.019) | 1.289 (0.114) |
| Super Enhancer + 500bp | 0.172 | 0.251 (0.022) | 1.464 (0.129) |
| TFBS | 0.132 | 0.357 (0.087) | 2.692 (0.659) |
| TFBS + 500bp | 0.343 | 0.438 (0.071) | 1.275 (0.207) |
| Transcribed | 0.345 | 0.442 (0.088) | 1.280 (0.256) |
| Transcribed + 500bp | 0.763 | 0.775 (0.051) | 1.015 (0.067) |
| TSS | 0.018 | 0.059 (0.030) | 3.239 (1.670) |
| TSS + 500bp | 0.035 | 0.097 (0.030) | 2.792 (0.874) |
| 3-prime UTR | 0.011 | 0.059 (0.022) | 5.352 (2.007) |
| 3-prime UTR + 500bp | 0.027 | 0.054 (0.021) | 2.013 (0.796) |
| 5-prime UTR | 0.005 | 0.035 (0.018) | 6.541 (3.287) |
| 5-prime UTR + 500bp | 0.028 | 0.055 (0.022) | 1.984 (0.794) |
| Weak Enhancer | 0.021 | 0.119 (0.039) | 5.640 (1.837) |
| Weak Enhancer + 500bp | 0.089 | 0.236 (0.044) | 2.658 (0.495) |

**(S4.C).** Proportion of heritability and enrichment for different functional categories for Age at menarche.

(S4.B) Proportion of heritability and enrichment for different functional categories for BMI.
| Annotation | Prop. SNPs | Prop. $h^2$ | Enrichment |
| --- | --- | --- | --- |
| Coding | 0.015 | 0.123 (0.030) | 8.413 (2.059) |
| Coding + 500bp | 0.065 | 0.094 (0.035) | 1.453 (0.537) |
| Conserved | 0.026 | 0.462 (0.063) | 17.725 (2.414) |
| Conserved + 500bp | 0.333 | 0.616 (0.090) | 1.853 (0.271) |
| CTCF | 0.024 | 0.022 (0.051) | 0.935 (2.150) |
| CTCF + 500bp | 0.071 | 0.049 (0.053) | 0.689 (0.749) |
| DGF | 0.138 | 0.207 (0.113) | 1.508 (0.823) |
| DGF + 500bp | 0.542 | 0.429 (0.084) | 0.793 (0.155) |
| DHS peaks | 0.112 | 0.008 (0.120) | 0.071 (1.071) |
| DHS | 0.168 | 0.052 (0.135) | 0.311 (0.806) |
| DHS + 500bp | 0.499 | 0.631 (0.096) | 1.265 (0.193) |
| FANTOM5 Enhancer | 0.004 | -0.000 (0.022) | -0.028 (5.150) |
| FANTOM5 Enhancer + 500bp | 0.019 | 0.006 (0.025) | 0.321 (1.315) |
| Enhancer | 0.063 | 0.137 (0.061) | 2.163 (0.960) |
| Enhancer + 500bp | 0.154 | 0.215 (0.059) | 1.397 (0.384) |
| Fetal DHS | 0.085 | 0.108 (0.093) | 1.272 (1.100) |
| Fetal DHS + 500bp | 0.285 | 0.291 (0.101) | 1.020 (0.356) |
| H3K27ac<sup>19</sup> | 0.391 | 0.455 (0.037) | 1.164 (0.094) |
| H3K27ac<sup>19</sup> + 500bp | 0.423 | 0.489 (0.052) | 1.158 (0.123) |
| H3K27ac<sup>18</sup> | 0.269 | 0.409 (0.071) | 1.517 (0.264) |
| H3K27ac<sup>18</sup> + 500bp | 0.336 | 0.486 (0.070) | 1.446 (0.207) |
| H3K4me1 peaks | 0.171 | 0.471 (0.116) | 2.747 (0.678) |
| H3K4me1 | 0.427 | 0.621 (0.085) | 1.455 (0.200) |
| H3K4me1 + 500bp | 0.609 | 0.847 (0.057) | 1.391 (0.094) |
| H3K4me3 peaks | 0.042 | 0.211 (0.071) | 5.042 (1.695) |
| H3K4me3 | 0.133 | 0.237 (0.067) | 1.779 (0.502) |
| H3K4me3 + 500bp | 0.255 | 0.308 (0.064) | 1.207 (0.249) |
| H3K9ac peaks | 0.039 | 0.220 (0.062) | 5.671 (1.596) |
| H3K9ac | 0.126 | 0.345 (0.066) | 2.736 (0.520) |
| H3K9ac + 500bp | 0.231 | 0.460 (0.055) | 1.996 (0.240) |
| Intron | 0.387 | 0.437 (0.035) | 1.129 (0.090) |
| Intron + 500bp | 0.397 | 0.537 (0.031) | 1.353 (0.078) |
| PromoterFlanking | 0.008 | 0.020 (0.031) | 2.394 (3.660) |
| PromoterFlanking + 500bp | 0.033 | 0.056 (0.033) | 1.684 (0.992) |
| Promoter | 0.031 | 0.123 (0.033) | 3.945 (1.051) |
| Promoter + 500bp | 0.039 | 0.116 (0.025) | 2.995 (0.660) |
| Repressed | 0.461 | 0.385 (0.100) | 0.835 (0.216) |
| Repressed + 500bp | 0.719 | 0.513 (0.038) | 0.714 (0.053) |
| Super Enhancer | 0.168 | 0.213 (0.030) | 1.264 (0.176) |
| Super Enhancer + 500bp | 0.172 | 0.233 (0.029) | 1.357 (0.171) |
| TFBS | 0.132 | 0.437 (0.098) | 3.301 (0.737) |
| TFBS + 500bp | 0.343 | 0.427 (0.092) | 1.244 (0.268) |
| Transcribed | 0.345 | 0.436 (0.086) | 1.263 (0.250) |
| Transcribed + 500bp | 0.763 | 0.681 (0.057) | 0.893 (0.075) |
| TSS | 0.018 | 0.165 (0.043) | 9.064 (2.354) |
| TSS + 500bp | 0.035 | 0.167 (0.036) | 4.786 (1.040) |
| 3-prime UTR | 0.011 | 0.034 (0.022) | 3.053 (1.948) |
| 3-prime UTR + 500bp | 0.027 | 0.047 (0.020) | 1.762 (0.761) |
| 5-prime UTR | 0.005 | 0.042 (0.021) | 7.792 (3.826) |
| 5-prime UTR + 500bp | 0.028 | 0.059 (0.031) | 2.129 (1.101) |
| Weak Enhancer | 0.021 | 0.011 (0.045) | 0.542 (2.148) |
| Weak Enhancer + 500bp | 0.089 | 0.113 (0.054) | 1.274 (0.602) |

**(S4.D).** Proportion of heritability and enrichment for different functional categories for LDL.

(S4.C) Proportion of heritability and enrichment for different functional categories for Age at menarche.
| Annotation | Prop. SNPs | Prop. $h^2$ | Enrichment |
| --- | --- | --- | --- |
| Coding | 0.015 | 0.128 (0.093) | 8.740 (6.337) |
| Coding + 500bp | 0.065 | 0.502 (0.089) | 7.782 (1.380) |
| Conserved | 0.026 | 0.264 (0.124) | 10.111 (4.756) |
| Conserved + 500bp | 0.333 | 0.811 (0.143) | 2.438 (0.429) |
| CTCF | 0.024 | 0.179 (0.109) | 7.506 (4.590) |
| CTCF + 500bp | 0.071 | 0.079 (0.106) | 1.109 (1.493) |
| DGF | 0.138 | 0.542 (0.192) | 3.938 (1.398) |
| DGF + 500bp | 0.542 | 0.806 (0.149) | 1.489 (0.276) |
| DHS peaks | 0.112 | 0.417 (0.189) | 3.730 (1.691) |
| DHS | 0.168 | 0.505 (0.247) | 3.008 (1.473) |
| DHS + 500bp | 0.499 | 0.810 (0.162) | 1.623 (0.325) |
| FANTOM5 Enhancer | 0.004 | -0.050 (0.047) | -11.493 (10.749) |
| FANTOM5 Enhancer + 500bp | 0.019 | -0.006 (0.047) | -0.323 (2.442) |
| Enhancer | 0.063 | 0.500 (0.157) | 7.895 (2.477) |
| Enhancer + 500bp | 0.154 | 0.524 (0.121) | 3.406 (0.783) |
| Fetal DHS | 0.085 | 0.370 (0.194) | 4.370 (2.290) |
| Fetal DHS + 500bp | 0.285 | 0.783 (0.133) | 2.746 (0.466) |
| H3K27ac<sup>19</sup> | 0.391 | 0.776 (0.093) | 1.984 (0.237) |
| H3K27ac<sup>19</sup> + 500bp | 0.423 | 0.962 (0.075) | 2.277 (0.177) |
| H3K27ac<sup>18</sup> | 0.269 | 0.829 (0.155) | 3.075 (0.575) |
| H3K27ac<sup>18</sup> + 500bp | 0.336 | 0.772 (0.100) | 2.298 (0.298) |
| H3K4me1 peaks | 0.171 | 0.430 (0.264) | 2.512 (1.539) |
| H3K4me1 | 0.427 | 1.179 (0.151) | 2.765 (0.354) |
| H3K4me1 + 500bp | 0.609 | 1.056 (0.094) | 1.733 (0.154) |
| H3K4me3 peaks | 0.042 | 0.069 (0.186) | 1.658 (4.459) |
| H3K4me3 | 0.133 | 0.628 (0.173) | 4.715 (1.296) |
| H3K4me3 + 500bp | 0.255 | 0.933 (0.181) | 3.652 (0.710) |
| H3K9ac peaks | 0.039 | -0.028 (0.165) | -0.712 (4.249) |
| H3K9ac | 0.126 | 0.751 (0.167) | 5.951 (1.324) |
| H3K9ac + 500bp | 0.231 | 0.832 (0.187) | 3.607 (0.812) |
| Intron | 0.387 | 0.472 (0.082) | 1.218 (0.212) |
| Intron + 500bp | 0.397 | 0.657 (0.050) | 1.654 (0.126) |
| PromoterFlanking | 0.008 | 0.055 (0.073) | 6.544 (8.609) |
| PromoterFlanking + 500bp | 0.033 | 0.264 (0.087) | 7.895 (2.605) |
| Promoter | 0.031 | 0.140 (0.073) | 4.499 (2.352) |
| Promoter + 500bp | 0.039 | 0.173 (0.048) | 4.475 (1.248) |
| Repressed | 0.461 | -0.077 (0.140) | -0.167 (0.304) |
| Repressed + 500bp | 0.719 | 0.168 (0.077) | 0.234 (0.107) |
| Super Enhancer | 0.168 | 0.439 (0.059) | 2.608 (0.351) |
| Super Enhancer + 500bp | 0.172 | 0.467 (0.059) | 2.721 (0.345) |
| TFBS | 0.132 | 0.871 (0.231) | 6.576 (1.747) |
| TFBS + 500bp | 0.343 | 0.787 (0.152) | 2.291 (0.443) |
| Transcribed | 0.345 | 0.573 (0.139) | 1.659 (0.403) |
| Transcribed + 500bp | 0.763 | 0.627 (0.106) | 0.821 (0.139) |
| TSS | 0.018 | 0.086 (0.097) | 4.746 (5.306) |
| TSS + 500bp | 0.035 | 0.452 (0.083) | 12.976 (2.387) |
| 3-prime UTR | 0.011 | 0.035 (0.045) | 3.190 (4.086) |
| 3-prime UTR + 500bp | 0.027 | 0.109 (0.041) | 4.044 (1.528) |
| 5-prime UTR | 0.005 | 0.015 (0.041) | 2.768 (7.486) |
| 5-prime UTR + 500bp | 0.028 | 0.088 (0.069) | 3.154 (2.471) |
| Weak Enhancer | 0.021 | 0.145 (0.096) | 6.876 (4.532) |
| Weak Enhancer + 500bp | 0.089 | 0.257 (0.109) | 2.890 (1.221) |

**(S4.E).** Proportion of heritability and enrichment for different functional categories for HDL.

(S4.D) Proportion of heritability and enrichment for different functional categories for LDL.
| Annotation | Prop. SNPs | Prop. $h^2$ | Enrichment |
| --- | --- | --- | --- |
| Coding | 0.015 | 0.058 (0.058) | 3.986 (3.937) |
| Coding + 500bp | 0.065 | 0.309 (0.064) | 4.788 (0.993) |
| Conserved | 0.026 | 0.269 (0.075) | 10.339 (2.870) |
| Conserved + 500bp | 0.333 | 0.582 (0.121) | 1.749 (0.364) |
| CTCF | 0.024 | -0.049 (0.068) | -2.077 (2.851) |
| CTCF + 500bp | 0.071 | 0.032 (0.084) | 0.453 (1.178) |
| DGF | 0.138 | 0.093 (0.195) | 0.676 (1.420) |
| DGF + 500bp | 0.542 | 0.735 (0.149) | 1.358 (0.275) |
| DHS peaks | 0.112 | 0.219 (0.193) | 1.960 (1.727) |
| DHS | 0.168 | 0.289 (0.190) | 1.721 (1.132) |
| DHS + 500bp | 0.499 | 0.921 (0.214) | 1.846 (0.428) |
| FANTOM5 Enhancer | 0.004 | 0.001 (0.049) | 0.162 (11.325) |
| FANTOM5 Enhancer + 500bp | 0.019 | -0.004 (0.051) | -0.213 (2.700) |
| Enhancer | 0.063 | 0.211 (0.125) | 3.332 (1.971) |
| Enhancer + 500bp | 0.154 | 0.512 (0.101) | 3.327 (0.658) |
| Fetal DHS | 0.085 | 0.091 (0.137) | 1.079 (1.620) |
| Fetal DHS + 500bp | 0.285 | 0.736 (0.116) | 2.583 (0.407) |
| H3K27ac<sup>19</sup> | 0.391 | 0.802 (0.067) | 2.051 (0.172) |
| H3K27ac<sup>19</sup> + 500bp | 0.423 | 0.828 (0.072) | 1.960 (0.171) |
| H3K27ac<sup>18</sup> | 0.269 | 0.904 (0.190) | 3.353 (0.705) |
| H3K27ac<sup>18</sup> + 500bp | 0.336 | 0.839 (0.082) | 2.495 (0.244) |
| H3K4me1 peaks | 0.171 | 0.602 (0.221) | 3.515 (1.291) |
| H3K4me1 | 0.427 | 1.055 (0.165) | 2.474 (0.388) |
| H3K4me1 + 500bp | 0.609 | 1.035 (0.093) | 1.699 (0.153) |
| H3K4me3 peaks | 0.042 | 0.019 (0.129) | 0.451 (3.082) |
| H3K4me3 | 0.133 | 0.530 (0.129) | 3.975 (0.970) |
| H3K4me3 + 500bp | 0.255 | 0.690 (0.111) | 2.702 (0.435) |
| H3K9ac peaks | 0.039 | 0.240 (0.121) | 6.195 (3.118) |
| H3K9ac | 0.126 | 0.632 (0.106) | 5.015 (0.842) |
| H3K9ac + 500bp | 0.231 | 0.856 (0.110) | 3.714 (0.477) |
| Intron | 0.387 | 0.523 (0.070) | 1.349 (0.180) |
| Intron + 500bp | 0.397 | 0.603 (0.048) | 1.518 (0.120) |
| PromoterFlanking | 0.008 | -0.039 (0.038) | -4.590 (4.512) |
| PromoterFlanking + 500bp | 0.033 | 0.071 (0.058) | 2.111 (1.727) |
| Promoter | 0.031 | 0.180 (0.073) | 5.785 (2.340) |
| Promoter + 500bp | 0.039 | 0.162 (0.046) | 4.197 (1.182) |
| Repressed | 0.461 | 0.143 (0.114) | 0.310 (0.246) |
| Repressed + 500bp | 0.719 | 0.312 (0.072) | 0.434 (0.100) |
| Super Enhancer | 0.168 | 0.531 (0.051) | 3.151 (0.305) |
| Super Enhancer + 500bp | 0.172 | 0.535 (0.048) | 3.120 (0.281) |
| TFBS | 0.132 | 0.358 (0.189) | 2.706 (1.428) |
| TFBS + 500bp | 0.343 | 0.601 (0.142) | 1.749 (0.413) |
| Transcribed | 0.345 | 0.445 (0.108) | 1.289 (0.314) |
| Transcribed + 500bp | 0.763 | 0.839 (0.070) | 1.100 (0.091) |
| TSS | 0.018 | 0.066 (0.057) | 3.601 (3.136) |
| TSS + 500bp | 0.035 | 0.306 (0.072) | 8.782 (2.072) |
| 3-prime UTR | 0.011 | -0.014 (0.036) | -1.224 (3.289) |
| 3-prime UTR + 500bp | 0.027 | 0.071 (0.040) | 2.623 (1.498) |
| 5-prime UTR | 0.005 | 0.082 (0.041) | 15.172 (7.520) |
| 5-prime UTR + 500bp | 0.028 | 0.086 (0.039) | 3.076 (1.410) |
| Weak Enhancer | 0.021 | 0.081 (0.077) | 3.859 (3.629) |
| Weak Enhancer + 500bp | 0.089 | 0.321 (0.105) | 3.605 (1.181) |

**(S4.F).**
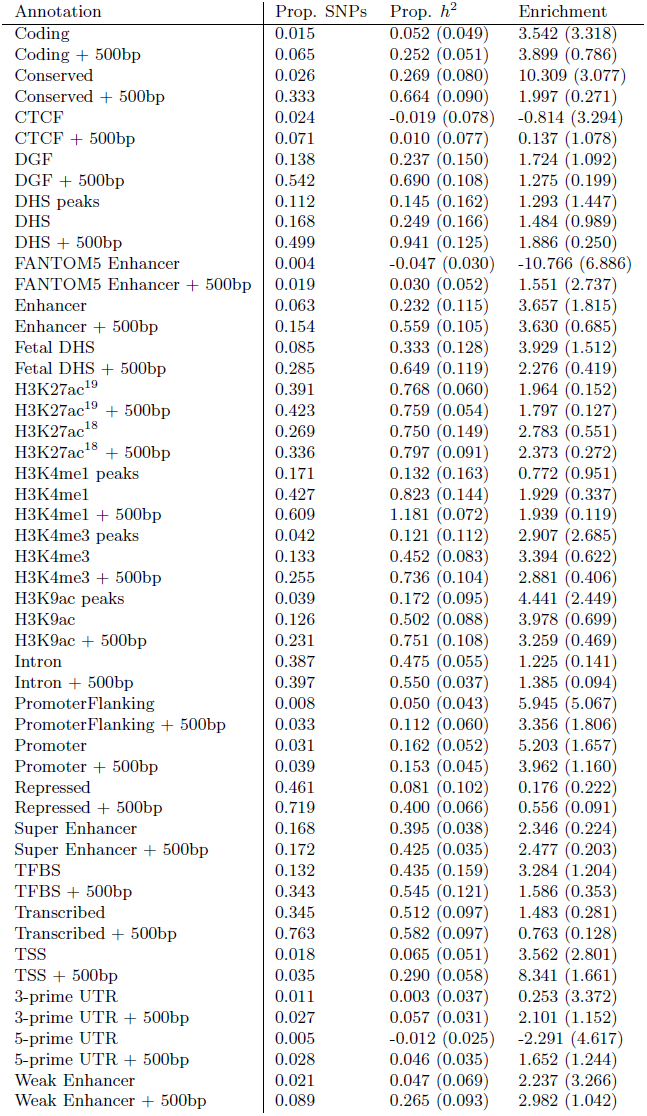
Proportion of heritability and enrichment for different functional categories for Triglycerides.

**(S4.G).** Proportion of heritability and enrichment for different functional categories for Coronary artery disease.

(S4.F) Proportion of heritability and enrichment for different functional categories for Triglycerides.
| Annotation | Prop. SNPs | Prop. $h^2$ | Enrichment |
| --- | --- | --- | --- |
| Coding | 0.015 | 0.074 (0.057) | 5.050 (3.886) |
| Coding + 500bp | 0.065 | 0.261 (0.068) | 4.043 (1.049) |
| Conserved | 0.026 | 0.203 (0.096) | 7.808 (3.677) |
| Conserved + 500bp | 0.333 | 0.568 (0.144) | 1.708 (0.433) |
| CTCF | 0.024 | -0.040 (0.105) | -1.659 (4.397) |
| CTCF + 500bp | 0.071 | -0.070 (0.122) | -0.980 (1.713) |
| DGF | 0.138 | 0.743 (0.256) | 5.402 (1.858) |
| DGF + 500bp | 0.542 | 0.860 (0.143) | 1.587 (0.265) |
| DHS peaks | 0.112 | 0.361 (0.254) | 3.229 (2.271) |
| DHS | 0.168 | 0.514 (0.264) | 3.063 (1.575) |
| DHS + 500bp | 0.499 | 0.943 (0.164) | 1.891 (0.330) |
| FANTOM5 Enhancer | 0.004 | -0.040 (0.053) | -9.331 (12.124) |
| FANTOM5 Enhancer + 500bp | 0.019 | 0.045 (0.063) | 2.359 (3.329) |
| Enhancer | 0.063 | 0.394 (0.130) | 6.217 (2.046) |
| Enhancer + 500bp | 0.154 | 0.406 (0.120) | 2.635 (0.777) |
| Fetal DHS | 0.085 | 0.282 (0.200) | 3.324 (2.364) |
| Fetal DHS + 500bp | 0.285 | 0.746 (0.164) | 2.618 (0.575) |
| H3K27ac<sup>19</sup> | 0.391 | 0.727 (0.087) | 1.859 (0.221) |
| H3K27ac<sup>19</sup> + 500bp | 0.423 | 0.801 (0.084) | 1.895 (0.198) |
| H3K27ac<sup>18</sup> | 0.269 | 0.639 (0.141) | 2.370 (0.523) |
| H3K27ac<sup>18</sup> + 500bp | 0.336 | 0.837 (0.129) | 2.490 (0.384) |
| H3K4me1 peaks | 0.171 | 0.497 (0.197) | 2.901 (1.148) |
| H3K4me1 | 0.427 | 0.783 (0.164) | 1.835 (0.384) |
| H3K4me1 + 500bp | 0.609 | 1.137 (0.086) | 1.866 (0.141) |
| H3K4me3 peaks | 0.042 | 0.266 (0.156) | 6.363 (3.724) |
| H3K4me3 | 0.133 | 0.288 (0.126) | 2.161 (0.949) |
| H3K4me3 + 500bp | 0.255 | 0.890 (0.146) | 3.482 (0.571) |
| H3K9ac peaks | 0.039 | 0.381 (0.141) | 9.820 (3.633) |
| H3K9ac | 0.126 | 0.686 (0.139) | 5.439 (1.105) |
| H3K9ac + 500bp | 0.231 | 0.788 (0.123) | 3.418 (0.535) |
| Intron | 0.387 | 0.483 (0.065) | 1.248 (0.167) |
| Intron + 500bp | 0.397 | 0.525 (0.049) | 1.323 (0.124) |
| PromoterFlanking | 0.008 | 0.001 (0.057) | 0.071 (6.744) |
| PromoterFlanking + 500bp | 0.033 | 0.108 (0.071) | 3.237 (2.115) |
| Promoter | 0.031 | 0.104 (0.071) | 3.344 (2.270) |
| Promoter + 500bp | 0.039 | 0.068 (0.047) | 1.765 (1.220) |
| Repressed | 0.461 | 0.395 (0.168) | 0.856 (0.365) |
| Repressed + 500bp | 0.719 | 0.359 (0.074) | 0.499 (0.103) |
| Super Enhancer | 0.168 | 0.414 (0.053) | 2.459 (0.317) |
| Super Enhancer + 500bp | 0.172 | 0.415 (0.058) | 2.418 (0.337) |
| TFBS | 0.132 | 0.364 (0.211) | 2.751 (1.595) |
| TFBS + 500bp | 0.343 | 0.487 (0.156) | 1.418 (0.454) |
| Transcribed | 0.345 | 0.302 (0.142) | 0.874 (0.410) |
| Transcribed + 500bp | 0.763 | 0.677 (0.113) | 0.887 (0.148) |
| TSS | 0.018 | 0.185 (0.077) | 10.152 (4.239) |
| TSS + 500bp | 0.035 | 0.141 (0.080) | 4.061 (2.291) |
| 3-prime UTR | 0.011 | 0.055 (0.045) | 5.018 (4.040) |
| 3-prime UTR + 500bp | 0.027 | 0.151 (0.064) | 5.624 (2.358) |
| 5-prime UTR | 0.005 | -0.003 (0.037) | -0.542 (6.731) |
| 5-prime UTR + 500bp | 0.028 | 0.029 (0.054) | 1.036 (1.958) |
| Weak Enhancer | 0.021 | 0.128 (0.091) | 6.055 (4.326) |
| Weak Enhancer + 500bp | 0.089 | 0.285 (0.113) | 3.200 (1.266) |

**(S4.H).**
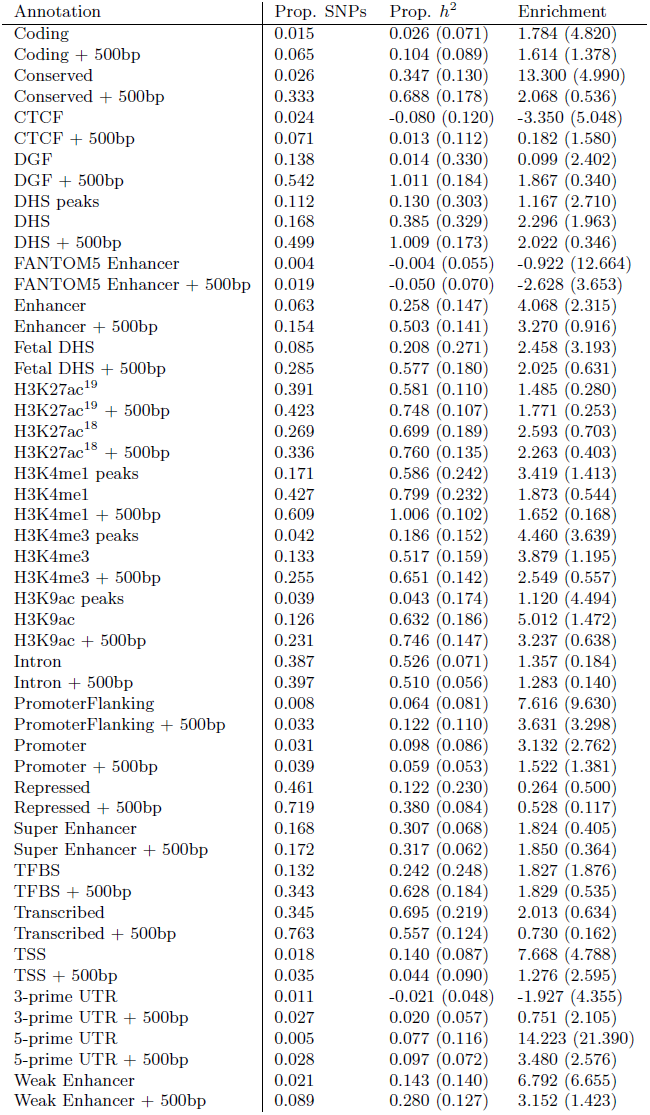
Proportion of heritability and enrichment for different functional categories for Type 2 Diabetes.

(S4.G) Proportion of heritability and enrichment for different functional categories for Coronary artery disease.
| Annotation | Prop. SNPs | Prop. $h^2$ | Enrichment |
| --- | --- | --- | --- |
| Coding | 0.015 | 0.026 (0.071) | 1.784 (4.820) |
| Coding + 500bp | 0.065 | 0.104 (0.089) | 1.614 (1.378) |
| Conserved | 0.026 | 0.347 (0.130) | 13.300 (4.990) |
| Conserved + 500bp | 0.333 | 0.688 (0.178) | 2.068 (0.536) |
| CTCF | 0.024 | -0.080 (0.120) | -3.350 (5.048) |
| CTCF + 500bp | 0.071 | 0.013 (0.112) | 0.182 (1.580) |
| DGF | 0.138 | 0.014 (0.330) | 0.099 (2.402) |
| DGF + 500bp | 0.542 | 1.011 (0.184) | 1.867 (0.340) |
| DHS peaks | 0.112 | 0.130 (0.303) | 1.167 (2.710) |
| DHS | 0.168 | 0.385 (0.329) | 2.296 (1.963) |
| DHS + 500bp | 0.499 | 1.009 (0.173) | 2.022 (0.346) |
| FANTOM5 Enhancer | 0.004 | -0.004 (0.055) | -0.922 (12.664) |
| FANTOM5 Enhancer + 500bp | 0.019 | -0.050 (0.070) | -2.628 (3.653) |
| Enhancer | 0.063 | 0.258 (0.147) | 4.068 (2.315) |
| Enhancer + 500bp | 0.154 | 0.503 (0.141) | 3.270 (0.916) |
| Fetal DHS | 0.085 | 0.208 (0.271) | 2.458 (3.193) |
| Fetal DHS + 500bp | 0.285 | 0.577 (0.180) | 2.025 (0.631) |
| H3K27ac<sup>19</sup> | 0.391 | 0.581 (0.110) | 1.485 (0.280) |
| H3K27ac<sup>19</sup> + 500bp | 0.423 | 0.748 (0.107) | 1.771 (0.253) |
| H3K27ac<sup>18</sup> | 0.269 | 0.699 (0.189) | 2.593 (0.703) |
| H3K27ac<sup>18</sup> + 500bp | 0.336 | 0.760 (0.135) | 2.263 (0.403) |
| H3K4me1 peaks | 0.171 | 0.586 (0.242) | 3.419 (1.413) |
| H3K4me1 | 0.427 | 0.799 (0.232) | 1.873 (0.544) |
| H3K4me1 + 500bp | 0.609 | 1.006 (0.102) | 1.652 (0.168) |
| H3K4me3 peaks | 0.042 | 0.186 (0.152) | 4.460 (3.639) |
| H3K4me3 | 0.133 | 0.517 (0.159) | 3.879 (1.195) |
| H3K4me3 + 500bp | 0.255 | 0.651 (0.142) | 2.549 (0.557) |
| H3K9ac peaks | 0.039 | 0.043 (0.174) | 1.120 (4.494) |
| H3K9ac | 0.126 | 0.632 (0.186) | 5.012 (1.472) |
| H3K9ac + 500bp | 0.231 | 0.746 (0.147) | 3.237 (0.638) |
| Intron | 0.387 | 0.526 (0.071) | 1.357 (0.184) |
| Intron + 500bp | 0.397 | 0.510 (0.056) | 1.283 (0.140) |
| PromoterFlanking | 0.008 | 0.064 (0.081) | 7.616 (9.630) |
| PromoterFlanking + 500bp | 0.033 | 0.122 (0.110) | 3.631 (3.298) |
| Promoter | 0.031 | 0.098 (0.086) | 3.132 (2.762) |
| Promoter + 500bp | 0.039 | 0.059 (0.053) | 1.522 (1.381) |
| Repressed | 0.461 | 0.122 (0.230) | 0.264 (0.500) |
| Repressed + 500bp | 0.719 | 0.380 (0.084) | 0.528 (0.117) |
| Super Enhancer | 0.168 | 0.307 (0.068) | 1.824 (0.405) |
| Super Enhancer + 500bp | 0.172 | 0.317 (0.062) | 1.850 (0.364) |
| TFBS | 0.132 | 0.242 (0.248) | 1.827 (1.876) |
| TFBS + 500bp | 0.343 | 0.628 (0.184) | 1.829 (0.535) |
| Transcribed | 0.345 | 0.695 (0.219) | 2.013 (0.634) |
| Transcribed + 500bp | 0.763 | 0.557 (0.124) | 0.730 (0.162) |
| TSS | 0.018 | 0.140 (0.087) | 7.668 (4.788) |
| TSS + 500bp | 0.035 | 0.044 (0.090) | 1.276 (2.595) |
| 3-prime UTR | 0.011 | -0.021 (0.048) | -1.927 (4.355) |
| 3-prime UTR + 500bp | 0.027 | 0.020 (0.057) | 0.751 (2.105) |
| 5-prime UTR | 0.005 | 0.077 (0.116) | 14.223 (21.390) |
| 5-prime UTR + 500bp | 0.028 | 0.097 (0.072) | 3.480 (2.576) |
| Weak Enhancer | 0.021 | 0.143 (0.140) | 6.792 (6.655) |
| Weak Enhancer + 500bp | 0.089 | 0.280 (0.127) | 3.152 (1.423) |

**(S4.I).** Proportion of heritability and enrichment for different functional categories for Fasting Glucose.

(S4.H) Proportion of heritability and enrichment for different functional categories for Type 2 Diabetes.
| Annotation | Prop. SNPs | Prop. $h^2$ | Enrichment |
| --- | --- | --- | --- |
| Coding | 0.015 | -0.014 (0.085) | -0.960 (5.787) |
| Coding + 500bp | 0.065 | 0.206 (0.087) | 3.184 (1.342) |
| Conserved | 0.026 | 0.286 (0.120) | 10.982 (4.588) |
| Conserved + 500bp | 0.333 | 0.874 (0.153) | 2.629 (0.460) |
| CTCF | 0.024 | -0.044 (0.117) | -1.842 (4.905) |
| CTCF + 500bp | 0.071 | -0.036 (0.132) | -0.501 (1.862) |
| DGF | 0.138 | 0.394 (0.297) | 2.861 (2.156) |
| DGF + 500bp | 0.542 | 0.723 (0.173) | 1.335 (0.319) |
| DHS peaks | 0.112 | 0.170 (0.291) | 1.523 (2.607) |
| DHS | 0.168 | 0.711 (0.307) | 4.238 (1.829) |
| DHS + 500bp | 0.499 | 0.564 (0.204) | 1.132 (0.410) |
| FANTOM5 Enhancer | 0.004 | -0.001 (0.057) | -0.176 (13.183) |
| FANTOM5 Enhancer + 500bp | 0.019 | 0.057 (0.072) | 2.975 (3.775) |
| Enhancer | 0.063 | 0.358 (0.172) | 5.658 (2.710) |
| Enhancer + 500bp | 0.154 | 0.519 (0.139) | 3.370 (0.900) |
| Fetal DHS | 0.085 | 0.375 (0.215) | 4.424 (2.537) |
| Fetal DHS + 500bp | 0.285 | 0.763 (0.206) | 2.677 (0.722) |
| H3K27ac<sup>19</sup> | 0.391 | 0.679 (0.097) | 1.737 (0.248) |
| H3K27ac<sup>19</sup> + 500bp | 0.423 | 0.734 (0.096) | 1.737 (0.227) |
| H3K27ac<sup>18</sup> | 0.269 | 0.734 (0.175) | 2.723 (0.649) |
| H3K27ac<sup>18</sup> + 500bp | 0.336 | 0.952 (0.158) | 2.833 (0.471) |
| H3K4me1 peaks | 0.171 | 0.865 (0.281) | 5.049 (1.641) |
| H3K4me1 | 0.427 | 0.743 (0.179) | 1.742 (0.420) |
| H3K4me1 + 500bp | 0.609 | 1.017 (0.117) | 1.670 (0.193) |
| H3K4me3 peaks | 0.042 | -0.014 (0.157) | -0.323 (3.750) |
| H3K4me3 | 0.133 | 0.583 (0.154) | 4.372 (1.156) |
| H3K4me3 + 500bp | 0.255 | 0.638 (0.141) | 2.496 (0.553) |
| H3K9ac peaks | 0.039 | 0.297 (0.153) | 7.658 (3.941) |
| H3K9ac | 0.126 | 0.753 (0.173) | 5.973 (1.370) |
| H3K9ac + 500bp | 0.231 | 0.542 (0.151) | 2.350 (0.654) |
| Intron | 0.387 | 0.596 (0.086) | 1.539 (0.223) |
| Intron + 500bp | 0.397 | 0.593 (0.064) | 1.492 (0.162) |
| PromoterFlanking | 0.008 | -0.075 (0.071) | -8.868 (8.458) |
| PromoterFlanking + 500bp | 0.033 | 0.126 (0.084) | 3.750 (2.512) |
| Promoter | 0.031 | 0.087 (0.080) | 2.795 (2.576) |
| Promoter + 500bp | 0.039 | 0.163 (0.067) | 4.214 (1.722) |
| Repressed | 0.461 | 0.326 (0.195) | 0.707 (0.423) |
| Repressed + 500bp | 0.719 | 0.308 (0.096) | 0.429 (0.133) |
| Super Enhancer | 0.168 | 0.311 (0.064) | 1.848 (0.381) |
| Super Enhancer + 500bp | 0.172 | 0.436 (0.064) | 2.541 (0.371) |
| TFBS | 0.132 | 0.420 (0.230) | 3.173 (1.733) |
| TFBS + 500bp | 0.343 | 0.497 (0.179) | 1.446 (0.523) |
| Transcribed | 0.345 | 0.502 (0.186) | 1.454 (0.539) |
| Transcribed + 500bp | 0.763 | 0.511 (0.139) | 0.670 (0.182) |
| TSS | 0.018 | -0.066 (0.085) | -3.643 (4.645) |
| TSS + 500bp | 0.035 | 0.113 (0.093) | 3.243 (2.678) |
| 3-prime UTR | 0.011 | -0.019 (0.053) | -1.722 (4.823) |
| 3-prime UTR + 500bp | 0.027 | -0.010 (0.063) | -0.367 (2.325) |
| 5-prime UTR | 0.005 | 0.041 (0.084) | 7.573 (15.417) |
| 5-prime UTR + 500bp | 0.028 | 0.120 (0.066) | 4.312 (2.389) |
| Weak Enhancer | 0.021 | 0.210 (0.117) | 9.946 (5.568) |
| Weak Enhancer + 500bp | 0.089 | 0.313 (0.131) | 3.523 (1.472) |

**(S4.J).** Proportion of heritability and enrichment for different functional categories for Schizophrenia. For many annotations there is heritability in the 500bp anking regions, as cautioned by Gusev et al.^13^ We believe, as hypothesized by Gusev et al., that this anking heritability inates the estimates of DHS heritability in Gusev et al. However, our work confirms the main message of Gusev et al. that much of the heritability of many traits, including schizophrenia, is located in regulatory regions.

(S4.I) Proportion of heritability and enrichment for different functional categories for Fasting Glucose.
| Annotation | Prop. SNPs | Prop. $h^2$ | Enrichment |
| --- | --- | --- | --- |
| Coding | 0.015 | 0.085 (0.022) | 5.823 (1.489) |
| Coding + 500bp | 0.065 | 0.127 (0.026) | 1.970 (0.398) |
| Conserved | 0.026 | 0.287 (0.044) | 11.028 (1.677) |
| Conserved + 500bp | 0.333 | 0.658 (0.047) | 1.979 (0.141) |
| CTCF | 0.024 | -0.041 (0.032) | -1.716 (1.344) |
| CTCF + 500bp | 0.071 | 0.079 (0.033) | 1.118 (0.460) |
| DGF | 0.138 | 0.168 (0.083) | 1.222 (0.604) |
| DGF + 500bp | 0.542 | 0.667 (0.057) | 1.232 (0.106) |
| DHS peaks | 0.112 | 0.066 (0.070) | 0.587 (0.625) |
| DHS | 0.168 | 0.175 (0.083) | 1.045 (0.493) |
| DHS + 500bp | 0.499 | 0.687 (0.058) | 1.377 (0.115) |
| FANTOM5 Enhancer | 0.004 | -0.024 (0.015) | -5.438 (3.564) |
| FANTOM5 Enhancer + 500bp | 0.019 | -0.001 (0.020) | -0.068 (1.033) |
| Enhancer | 0.063 | 0.029 (0.043) | 0.465 (0.675) |
| Enhancer + 500bp | 0.154 | 0.238 (0.040) | 1.544 (0.263) |
| Fetal DHS | 0.085 | 0.216 (0.070) | 2.547 (0.825) |
| Fetal DHS + 500bp | 0.285 | 0.486 (0.060) | 1.705 (0.211) |
| H3K27ac<sup>19</sup> | 0.391 | 0.503 (0.027) | 1.287 (0.070) |
| H3K27ac<sup>19</sup> + 500bp | 0.423 | 0.552 (0.036) | 1.306 (0.085) |
| H3K27ac<sup>18</sup> | 0.269 | 0.372 (0.045) | 1.381 (0.168) |
| H3K27ac<sup>18</sup> + 500bp | 0.336 | 0.544 (0.042) | 1.618 (0.125) |
| H3K4me1 peaks | 0.171 | 0.412 (0.072) | 2.407 (0.420) |
| H3K4me1 | 0.427 | 0.646 (0.053) | 1.514 (0.124) |
| H3K4me1 + 500bp | 0.609 | 0.794 (0.034) | 1.304 (0.056) |
| H3K4me3 peaks | 0.042 | 0.188 (0.048) | 4.511 (1.144) |
| H3K4me3 | 0.133 | 0.292 (0.046) | 2.192 (0.342) |
| H3K4me3 + 500bp | 0.255 | 0.353 (0.040) | 1.383 (0.156) |
| H3K9ac peaks | 0.039 | 0.239 (0.042) | 6.163 (1.085) |
| H3K9ac | 0.126 | 0.287 (0.051) | 2.278 (0.404) |
| H3K9ac + 500bp | 0.231 | 0.457 (0.037) | 1.981 (0.161) |
| Intron | 0.387 | 0.513 (0.024) | 1.323 (0.063) |
| Intron + 500bp | 0.397 | 0.524 (0.019) | 1.321 (0.047) |
| PromoterFlanking | 0.008 | 0.029 (0.021) | 3.397 (2.545) |
| PromoterFlanking + 500bp | 0.033 | 0.070 (0.025) | 2.099 (0.754) |
| Promoter | 0.031 | 0.068 (0.028) | 2.192 (0.887) |
| Promoter + 500bp | 0.039 | 0.037 (0.021) | 0.952 (0.531) |
| Repressed | 0.461 | 0.450 (0.070) | 0.975 (0.152) |
| Repressed + 500bp | 0.719 | 0.627 (0.027) | 0.873 (0.038) |
| Super Enhancer | 0.168 | 0.247 (0.022) | 1.467 (0.130) |
| Super Enhancer + 500bp | 0.172 | 0.253 (0.019) | 1.475 (0.113) |
| TFBS | 0.132 | 0.217 (0.059) | 1.639 (0.447) |
| TFBS + 500bp | 0.343 | 0.372 (0.059) | 1.083 (0.171) |
| Transcribed | 0.345 | 0.391 (0.062) | 1.133 (0.180) |
| Transcribed + 500bp | 0.763 | 0.812 (0.035) | 1.064 (0.046) |
| TSS | 0.018 | 0.053 (0.027) | 2.918 (1.467) |
| TSS + 500bp | 0.035 | 0.107 (0.023) | 3.081 (0.662) |
| 3-prime UTR | 0.011 | 0.062 (0.019) | 5.583 (1.693) |
| 3-prime UTR + 500bp | 0.027 | 0.085 (0.020) | 3.140 (0.758) |
| 5-prime UTR | 0.005 | 0.020 (0.012) | 3.656 (2.215) |
| 5-prime UTR + 500bp | 0.028 | 0.055 (0.020) | 1.975 (0.710) |
| Weak Enhancer | 0.021 | 0.016 (0.032) | 0.759 (1.537) |
| Weak Enhancer + 500bp | 0.089 | 0.146 (0.040) | 1.643 (0.447) |

**(S4.K).** Proportion of heritability and enrichment for different functional categories for Bipolar disorder.

| Annotation | Prop. SNPs | Prop. $h^2$ | Enrichment |
| --- | --- | --- | --- |
| Coding | 0.015 | 0.186 (0.090) | 12.671 (6.108) |
| Coding + 500bp | 0.065 | 0.200 (0.086) | 3.098 (1.336) |
| Conserved | 0.026 | 0.302 (0.115) | 11.588 (4.413) |
| Conserved + 500bp | 0.333 | 0.527 (0.115) | 1.586 (0.345) |
| CTCF | 0.024 | 0.013 (0.111) | 0.556 (4.672) |
| CTCF + 500bp | 0.071 | 0.138 (0.104) | 1.944 (1.466) |
| DGF | 0.138 | 0.723 (0.241) | 5.251 (1.754) |
| DGF + 500bp | 0.542 | 0.854 (0.169) | 1.578 (0.313) |
| DHS peaks | 0.112 | 0.447 (0.211) | 4.001 (1.890) |
| DHS | 0.168 | 0.670 (0.251) | 3.993 (1.495) |
| DHS + 500bp | 0.499 | 0.735 (0.170) | 1.474 (0.340) |
| FANTOM5 Enhancer | 0.004 | 0.013 (0.046) | 2.987 (10.718) |
| FANTOM5 Enhancer + 500bp | 0.019 | -0.106 (0.053) | -5.564 (2.793) |
| Enhancer | 0.063 | 0.317 (0.130) | 4.999 (2.059) |
| Enhancer + 500bp | 0.154 | 0.299 (0.123) | 1.940 (0.797) |
| Fetal DHS | 0.085 | 0.264 (0.186) | 3.120 (2.199) |
| Fetal DHS + 500bp | 0.285 | 0.555 (0.169) | 1.947 (0.595) |
| H3K27ac<sup>19</sup> | 0.391 | 0.456 (0.089) | 1.166 (0.226) |
| H3K27ac<sup>19</sup> + 500bp | 0.423 | 0.564 (0.085) | 1.335 (0.202) |
| H3K27ac<sup>18</sup> | 0.269 | 0.574 (0.136) | 2.132 (0.505) |
| H3K27ac<sup>18</sup> + 500bp | 0.336 | 0.727 (0.116) | 2.162 (0.346) |
| H3K4me1 peaks | 0.171 | 0.719 (0.215) | 4.196 (1.256) |
| H3K4me1 | 0.427 | 0.970 (0.166) | 2.274 (0.389) |
| H3K4me1 + 500bp | 0.609 | 0.754 (0.099) | 1.238 (0.162) |
| H3K4me3 peaks | 0.042 | 0.038 (0.124) | 0.905 (2.967) |
| H3K4me3 | 0.133 | 0.573 (0.147) | 4.295 (1.101) |
| H3K4me3 + 500bp | 0.255 | 0.220 (0.121) | 0.861 (0.473) |
| H3K9ac peaks | 0.039 | 0.160 (0.125) | 4.124 (3.225) |
| H3K9ac | 0.126 | 0.450 (0.153) | 3.572 (1.214) |
| H3K9ac + 500bp | 0.231 | 0.502 (0.117) | 2.176 (0.508) |
| Intron | 0.387 | 0.509 (0.092) | 1.313 (0.237) |
| Intron + 500bp | 0.397 | 0.608 (0.066) | 1.531 (0.166) |
| PromoterFlanking | 0.008 | 0.051 (0.063) | 6.090 (7.507) |
| PromoterFlanking + 500bp | 0.033 | 0.081 (0.085) | 2.429 (2.541) |
| Promoter | 0.031 | 0.140 (0.085) | 4.500 (2.711) |
| Promoter + 500bp | 0.039 | 0.128 (0.065) | 3.317 (1.671) |
| Repressed | 0.461 | 0.402 (0.184) | 0.871 (0.400) |
| Repressed + 500bp | 0.719 | 0.460 (0.068) | 0.640 (0.094) |
| Super Enhancer | 0.168 | 0.288 (0.047) | 1.712 (0.278) |
| Super Enhancer + 500bp | 0.172 | 0.252 (0.043) | 1.469 (0.250) |
| TFBS | 0.132 | 0.459 (0.218) | 3.462 (1.645) |
| TFBS + 500bp | 0.343 | 0.507 (0.153) | 1.477 (0.444) |
| Transcribed | 0.345 | 0.311 (0.179) | 0.900 (0.518) |
| Transcribed + 500bp | 0.763 | 0.773 (0.107) | 1.013 (0.140) |
| TSS | 0.018 | 0.081 (0.070) | 4.435 (3.863) |
| TSS + 500bp | 0.035 | 0.244 (0.074) | 7.016 (2.135) |
| 3-prime UTR | 0.011 | 0.094 (0.062) | 8.508 (5.598) |
| 3-prime UTR + 500bp | 0.027 | 0.072 (0.052) | 2.664 (1.921) |
| 5-prime UTR | 0.005 | 0.122 (0.052) | 22.400 (9.543) |
| 5-prime UTR + 500bp | 0.028 | 0.129 (0.052) | 4.656 (1.873) |
| Weak Enhancer | 0.021 | 0.168 (0.103) | 7.988 (4.864) |
| Weak Enhancer + 500bp | 0.089 | 0.217 (0.098) | 2.435 (1.107) |

**(S4.L).** Proportion of heritability and enrichment for different functional categories for Anorexia.

(S4.K) Proportion of heritability and enrichment for different functional categories for Bipolar disorder.
| Annotation | Prop. SNPs | Prop. $h^2$ | Enrichment |
| --- | --- | --- | --- |
| Coding | 0.015 | 0.017 (0.068) | 1.145 (4.645) |
| Coding + 500bp | 0.065 | 0.140 (0.059) | 2.169 (0.915) |
| Conserved | 0.026 | 0.186 (0.110) | 7.128 (4.214) |
| Conserved + 500bp | 0.333 | 0.635 (0.124) | 1.910 (0.373) |
| CTCF | 0.024 | 0.019 (0.098) | 0.802 (4.130) |
| CTCF + 500bp | 0.071 | 0.012 (0.107) | 0.162 (1.499) |
| DGF | 0.138 | 0.257 (0.240) | 1.865 (1.743) |
| DGF + 500bp | 0.542 | 0.573 (0.144) | 1.058 (0.266) |
| DHS peaks | 0.112 | 0.041 (0.212) | 0.367 (1.894) |
| DHS | 0.168 | -0.021 (0.272) | -0.126 (1.623) |
| DHS + 500bp | 0.499 | 0.780 (0.173) | 1.564 (0.346) |
| FANTOM5 Enhancer | 0.004 | -0.118 (0.044) | -27.161 (10.087) |
| FANTOM5 Enhancer + 500bp | 0.019 | 0.037 (0.050) | 1.915 (2.600) |
| Enhancer | 0.063 | 0.117 (0.117) | 1.852 (1.845) |
| Enhancer + 500bp | 0.154 | 0.237 (0.112) | 1.537 (0.731) |
| Fetal DHS | 0.085 | 0.222 (0.199) | 2.618 (2.343) |
| Fetal DHS + 500bp | 0.285 | 0.458 (0.169) | 1.606 (0.595) |
| H3K27ac<sup>19</sup> | 0.391 | 0.528 (0.077) | 1.350 (0.197) |
| H3K27ac<sup>19</sup> + 500bp | 0.423 | 0.435 (0.089) | 1.030 (0.211) |
| H3K27ac<sup>18</sup> | 0.269 | 0.362 (0.136) | 1.343 (0.503) |
| H3K27ac<sup>18</sup> + 500bp | 0.336 | 0.467 (0.116) | 1.391 (0.344) |
| H3K4me1 peaks | 0.171 | 0.481 (0.215) | 2.806 (1.253) |
| H3K4me1 | 0.427 | 0.867 (0.153) | 2.033 (0.358) |
| H3K4me1 + 500bp | 0.609 | 0.798 (0.107) | 1.309 (0.175) |
| H3K4me3 peaks | 0.042 | 0.101 (0.116) | 2.424 (2.770) |
| H3K4me3 | 0.133 | 0.205 (0.124) | 1.538 (0.929) |
| H3K4me3 + 500bp | 0.255 | 0.277 (0.121) | 1.086 (0.472) |
| H3K9ac peaks | 0.039 | -0.025 (0.138) | -0.656 (3.572) |
| H3K9ac | 0.126 | 0.244 (0.129) | 1.935 (1.019) |
| H3K9ac + 500bp | 0.231 | 0.178 (0.122) | 0.771 (0.528) |
| Intron | 0.387 | 0.417 (0.066) | 1.076 (0.170) |
| Intron + 500bp | 0.397 | 0.429 (0.054) | 1.081 (0.137) |
| PromoterFlanking | 0.008 | 0.041 (0.060) | 4.904 (7.124) |
| PromoterFlanking + 500bp | 0.033 | 0.114 (0.077) | 3.412 (2.313) |
| Promoter | 0.031 | 0.022 (0.069) | 0.692 (2.223) |
| Promoter + 500bp | 0.039 | -0.027 (0.043) | -0.694 (1.116) |
| Repressed | 0.461 | 0.402 (0.179) | 0.871 (0.389) |
| Repressed + 500bp | 0.719 | 0.724 (0.063) | 1.006 (0.088) |
| Super Enhancer | 0.168 | 0.171 (0.048) | 1.017 (0.284) |
| Super Enhancer + 500bp | 0.172 | 0.238 (0.045) | 1.387 (0.263) |
| TFBS | 0.132 | 0.309 (0.177) | 2.330 (1.340) |
| TFBS + 500bp | 0.343 | 0.386 (0.159) | 1.125 (0.462) |
| Transcribed | 0.345 | 0.397 (0.158) | 1.150 (0.456) |
| Transcribed + 500bp | 0.763 | 0.685 (0.113) | 0.897 (0.147) |
| TSS | 0.018 | 0.078 (0.074) | 4.287 (4.044) |
| TSS + 500bp | 0.035 | 0.081 (0.062) | 2.336 (1.767) |
| 3-prime UTR | 0.011 | 0.005 (0.040) | 0.477 (3.639) |
| 3-prime UTR + 500bp | 0.027 | 0.052 (0.047) | 1.935 (1.737) |
| 5-prime UTR | 0.005 | 0.001 (0.039) | 0.261 (7.157) |
| 5-prime UTR + 500bp | 0.028 | 0.029 (0.053) | 1.041 (1.900) |
| Weak Enhancer | 0.021 | 0.126 (0.085) | 5.969 (4.039) |
| Weak Enhancer + 500bp | 0.089 | 0.139 (0.094) | 1.564 (1.058) |

**(S4.M).**
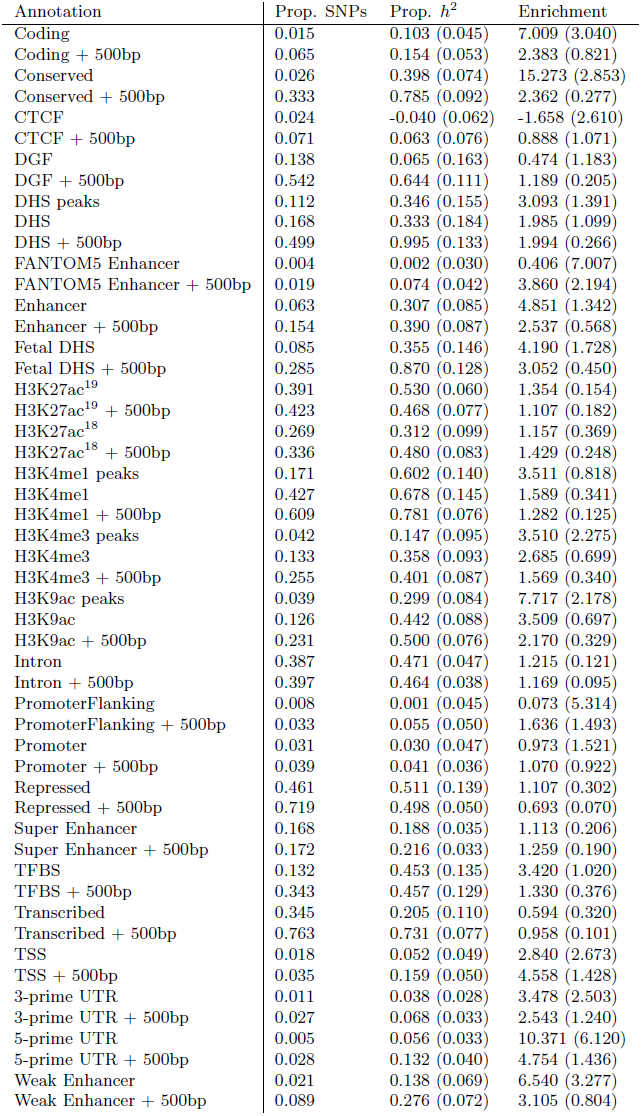
Proportion of heritability and enrichment for different functional categories for Years of education.

(S4.L) Proportion of heritability and enrichment for different functional categories for Anorexia.
| Annotation | Prop. SNPs | Prop. $h^2$ | Enrichment |
| --- | --- | --- | --- |
| Coding | 0.015 | 0.103 (0.045) | 7.009 (3.040) |
| Coding + 500bp | 0.065 | 0.154 (0.053) | 2.383 (0.821) |
| Conserved | 0.026 | 0.398 (0.074) | 15.273 (2.853) |
| Conserved + 500bp | 0.333 | 0.785 (0.092) | 2.362 (0.277) |
| CTCF | 0.024 | -0.040 (0.062) | -1.658 (2.610) |
| CTCF + 500bp | 0.071 | 0.063 (0.076) | 0.888 (1.071) |
| DGF | 0.138 | 0.065 (0.163) | 0.474 (1.183) |
| DGF + 500bp | 0.542 | 0.644 (0.111) | 1.189 (0.205) |
| DHS peaks | 0.112 | 0.346 (0.155) | 3.093 (1.391) |
| DHS | 0.168 | 0.333 (0.184) | 1.985 (1.099) |
| DHS + 500bp | 0.499 | 0.995 (0.133) | 1.994 (0.266) |
| FANTOM5 Enhancer | 0.004 | 0.002 (0.030) | 0.406 (7.007) |
| FANTOM5 Enhancer + 500bp | 0.019 | 0.074 (0.042) | 3.860 (2.194) |
| Enhancer | 0.063 | 0.307 (0.085) | 4.851 (1.342) |
| Enhancer + 500bp | 0.154 | 0.390 (0.087) | 2.537 (0.568) |
| Fetal DHS | 0.085 | 0.355 (0.146) | 4.190 (1.728) |
| Fetal DHS + 500bp | 0.285 | 0.870 (0.128) | 3.052 (0.450) |
| H3K27ac<sup>19</sup> | 0.391 | 0.530 (0.060) | 1.354 (0.154) |
| H3K27ac<sup>19</sup> + 500bp | 0.423 | 0.468 (0.077) | 1.107 (0.182) |
| H3K27ac<sup>18</sup> | 0.269 | 0.312 (0.099) | 1.157 (0.369) |
| H3K27ac<sup>18</sup> + 500bp | 0.336 | 0.480 (0.083) | 1.429 (0.248) |
| H3K4me1 peaks | 0.171 | 0.602 (0.140) | 3.511 (0.818) |
| H3K4me1 | 0.427 | 0.678 (0.145) | 1.589 (0.341) |
| H3K4me1 + 500bp | 0.609 | 0.781 (0.076) | 1.282 (0.125) |
| H3K4me3 peaks | 0.042 | 0.147 (0.095) | 3.510 (2.275) |
| H3K4me3 | 0.133 | 0.358 (0.093) | 2.685 (0.699) |
| H3K4me3 + 500bp | 0.255 | 0.401 (0.087) | 1.569 (0.340) |
| H3K9ac peaks | 0.039 | 0.299 (0.084) | 7.717 (2.178) |
| H3K9ac | 0.126 | 0.442 (0.088) | 3.509 (0.697) |
| H3K9ac + 500bp | 0.231 | 0.500 (0.076) | 2.170 (0.329) |
| Intron | 0.387 | 0.471 (0.047) | 1.215 (0.121) |
| Intron + 500bp | 0.397 | 0.464 (0.038) | 1.169 (0.095) |
| PromoterFlanking | 0.008 | 0.001 (0.045) | 0.073 (5.314) |
| PromoterFlanking + 500bp | 0.033 | 0.055 (0.050) | 1.636 (1.493) |
| Promoter | 0.031 | 0.030 (0.047) | 0.973 (1.521) |
| Promoter + 500bp | 0.039 | 0.041 (0.036) | 1.070 (0.922) |
| Repressed | 0.461 | 0.511 (0.139) | 1.107 (0.302) |
| Repressed + 500bp | 0.719 | 0.498 (0.050) | 0.693 (0.070) |
| Super Enhancer | 0.168 | 0.188 (0.035) | 1.113 (0.206) |
| Super Enhancer + 500bp | 0.172 | 0.216 (0.033) | 1.259 (0.190) |
| TFBS | 0.132 | 0.453 (0.135) | 3.420 (1.020) |
| TFBS + 500bp | 0.343 | 0.457 (0.129) | 1.330 (0.376) |
| Transcribed | 0.345 | 0.205 (0.110) | 0.594 (0.320) |
| Transcribed + 500bp | 0.763 | 0.731 (0.077) | 0.958 (0.101) |
| TSS | 0.018 | 0.052 (0.049) | 2.840 (2.673) |
| TSS + 500bp | 0.035 | 0.159 (0.050) | 4.558 (1.428) |
| 3-prime UTR | 0.011 | 0.038 (0.028) | 3.478 (2.503) |
| 3-prime UTR + 500bp | 0.027 | 0.068 (0.033) | 2.543 (1.240) |
| 5-prime UTR | 0.005 | 0.056 (0.033) | 10.371 (6.120) |
| 5-prime UTR + 500bp | 0.028 | 0.132 (0.040) | 4.754 (1.436) |
| Weak Enhancer | 0.021 | 0.138 (0.069) | 6.540 (3.277) |
| Weak Enhancer + 500bp | 0.089 | 0.276 (0.072) | 3.105 (0.804) |

**(S4.N).** Proportion of heritability and enrichment for different functional categories for Ever smoked.

(S4.M) Proportion of heritability and enrichment for different functional categories for Years of education.
| Annotation | Prop. SNPs | Prop. $h^2$ | Enrichment |
| --- | --- | --- | --- |
| Coding | 0.015 | 0.116 (0.077) | 7.929 (5.258) |
| Coding + 500bp | 0.065 | 0.114 (0.077) | 1.759 (1.188) |
| Conserved | 0.026 | 0.402 (0.125) | 15.414 (4.811) |
| Conserved + 500bp | 0.333 | 0.750 (0.134) | 2.256 (0.403) |
| CTCF | 0.024 | 0.095 (0.128) | 3.973 (5.371) |
| CTCF + 500bp | 0.071 | 0.215 (0.132) | 3.029 (1.862) |
| DGF | 0.138 | 0.419 (0.287) | 3.048 (2.088) |
| DGF + 500bp | 0.542 | 0.896 (0.208) | 1.655 (0.384) |
| DHS peaks | 0.112 | 0.352 (0.268) | 3.153 (2.396) |
| DHS | 0.168 | 0.433 (0.286) | 2.581 (1.706) |
| DHS + 500bp | 0.499 | 0.958 (0.206) | 1.921 (0.413) |
| FANTOM5 Enhancer | 0.004 | 0.006 (0.050) | 1.319 (11.432) |
| FANTOM5 Enhancer + 500bp | 0.019 | -0.109 (0.072) | -5.736 (3.778) |
| Enhancer | 0.063 | 0.326 (0.152) | 5.141 (2.408) |
| Enhancer + 500bp | 0.154 | 0.387 (0.166) | 2.513 (1.078) |
| Fetal DHS | 0.085 | 0.538 (0.241) | 6.346 (2.845) |
| Fetal DHS + 500bp | 0.285 | 0.808 (0.180) | 2.834 (0.630) |
| H3K27ac<sup>19</sup> | 0.391 | 0.494 (0.093) | 1.263 (0.237) |
| H3K27ac<sup>19</sup> + 500bp | 0.423 | 0.379 (0.131) | 0.897 (0.309) |
| H3K27ac<sup>18</sup> | 0.269 | 0.192 (0.178) | 0.713 (0.661) |
| H3K27ac<sup>18</sup> + 500bp | 0.336 | 0.549 (0.153) | 1.634 (0.455) |
| H3K4me1 peaks | 0.171 | 0.643 (0.257) | 3.751 (1.501) |
| H3K4me1 | 0.427 | 0.946 (0.224) | 2.218 (0.525) |
| H3K4me1 + 500bp | 0.609 | 0.750 (0.136) | 1.231 (0.223) |
| H3K4me3 peaks | 0.042 | 0.176 (0.180) | 4.210 (4.313) |
| H3K4me3 | 0.133 | 0.275 (0.154) | 2.066 (1.154) |
| H3K4me3 + 500bp | 0.255 | 0.259 (0.148) | 1.016 (0.581) |
| H3K9ac peaks | 0.039 | 0.233 (0.153) | 6.009 (3.951) |
| H3K9ac | 0.126 | 0.252 (0.151) | 2.002 (1.194) |
| H3K9ac + 500bp | 0.231 | 0.162 (0.147) | 0.702 (0.635) |
| Intron | 0.387 | 0.372 (0.081) | 0.960 (0.210) |
| Intron + 500bp | 0.397 | 0.443 (0.068) | 1.115 (0.171) |
| PromoterFlanking | 0.008 | -0.017 (0.061) | -1.974 (7.269) |
| PromoterFlanking + 500bp | 0.033 | -0.001 (0.094) | -0.032 (2.795) |
| Promoter | 0.031 | -0.010 (0.070) | -0.334 (2.245) |
| Promoter + 500bp | 0.039 | -0.019 (0.055) | -0.483 (1.420) |
| Repressed | 0.461 | 0.192 (0.218) | 0.417 (0.473) |
| Repressed + 500bp | 0.719 | 0.504 (0.094) | 0.701 (0.131) |
| Super Enhancer | 0.168 | 0.135 (0.061) | 0.800 (0.365) |
| Super Enhancer + 500bp | 0.172 | 0.071 (0.062) | 0.414 (0.359) |
| TFBS | 0.132 | 0.743 (0.245) | 5.613 (1.849) |
| TFBS + 500bp | 0.343 | 0.385 (0.217) | 1.121 (0.630) |
| Transcribed | 0.345 | 0.276 (0.197) | 0.798 (0.570) |
| Transcribed + 500bp | 0.763 | 0.739 (0.113) | 0.969 (0.148) |
| TSS | 0.018 | 0.065 (0.087) | 3.591 (4.777) |
| TSS + 500bp | 0.035 | 0.046 (0.081) | 1.309 (2.315) |
| 3-prime UTR | 0.011 | 0.009 (0.053) | 0.839 (4.758) |
| 3-prime UTR + 500bp | 0.027 | -0.005 (0.054) | -0.171 (2.012) |
| 5-prime UTR | 0.005 | -0.009 (0.046) | -1.695 (8.473) |
| 5-prime UTR + 500bp | 0.028 | 0.029 (0.059) | 1.048 (2.126) |
| Weak Enhancer | 0.021 | 0.136 (0.116) | 6.438 (5.507) |
| Weak Enhancer + 500bp | 0.089 | 0.212 (0.129) | 2.387 (1.450) |

**(S4.O).** Proportion of heritability and enrichment for different functional categories for Rheumatoid arthritis.

(S4.N) Proportion of heritability and enrichment for different functional categories for Ever smoked.
| Annotation | Prop. SNPs | Prop. $h^2$ | Enrichment |
| --- | --- | --- | --- |
| Coding | 0.015 | 0.174 (0.090) | 11.891 (6.155) |
| Coding + 500bp | 0.065 | 0.365 (0.081) | 5.659 (1.262) |
| Conserved | 0.026 | 0.109 (0.115) | 4.195 (4.402) |
| Conserved + 500bp | 0.333 | 0.771 (0.122) | 2.319 (0.368) |
| CTCF | 0.024 | -0.006 (0.113) | -0.261 (4.762) |
| CTCF + 500bp | 0.071 | -0.152 (0.114) | -2.140 (1.610) |
| DGF | 0.138 | 0.871 (0.251) | 6.331 (1.826) |
| DGF + 500bp | 0.542 | 1.065 (0.147) | 1.966 (0.271) |
| DHS peaks | 0.112 | 0.357 (0.255) | 3.195 (2.283) |
| DHS | 0.168 | 0.523 (0.272) | 3.121 (1.619) |
| DHS + 500bp | 0.499 | 0.858 (0.145) | 1.719 (0.290) |
| FANTOM5 Enhancer | 0.004 | 0.158 (0.071) | 36.410 (16.488) |
| FANTOM5 Enhancer + 500bp | 0.019 | 0.283 (0.084) | 14.848 (4.407) |
| Enhancer | 0.063 | 0.379 (0.154) | 5.984 (2.437) |
| Enhancer + 500bp | 0.154 | 0.668 (0.120) | 4.342 (0.778) |
| Fetal DHS | 0.085 | 0.567 (0.199) | 6.690 (2.345) |
| Fetal DHS + 500bp | 0.285 | 0.537 (0.157) | 1.882 (0.551) |
| H3K27ac<sup>19</sup> | 0.391 | 0.980 (0.086) | 2.506 (0.219) |
| H3K27ac<sup>19</sup> + 500bp | 0.423 | 0.957 (0.083) | 2.263 (0.195) |
| H3K27ac<sup>18</sup> | 0.269 | 0.951 (0.161) | 3.530 (0.597) |
| H3K27ac<sup>18</sup> + 500bp | 0.336 | 0.792 (0.112) | 2.358 (0.333) |
| H3K4me1 peaks | 0.171 | 0.502 (0.201) | 2.931 (1.173) |
| H3K4me1 | 0.427 | 0.884 (0.191) | 2.072 (0.448) |
| H3K4me1 + 500bp | 0.609 | 1.041 (0.085) | 1.709 (0.140) |
| H3K4me3 peaks | 0.042 | 0.313 (0.144) | 7.501 (3.435) |
| H3K4me3 | 0.133 | 0.750 (0.149) | 5.628 (1.119) |
| H3K4me3 + 500bp | 0.255 | 0.610 (0.124) | 2.387 (0.484) |
| H3K9ac peaks | 0.039 | 0.146 (0.139) | 3.770 (3.580) |
| H3K9ac | 0.126 | 0.335 (0.149) | 2.655 (1.178) |
| H3K9ac + 500bp | 0.231 | 0.434 (0.130) | 1.883 (0.562) |
| Intron | 0.387 | 0.416 (0.071) | 1.073 (0.184) |
| Intron + 500bp | 0.397 | 0.513 (0.052) | 1.291 (0.132) |
| PromoterFlanking | 0.008 | 0.017 (0.073) | 2.049 (8.639) |
| PromoterFlanking + 500bp | 0.033 | 0.228 (0.095) | 6.818 (2.846) |
| Promoter | 0.031 | 0.251 (0.079) | 8.046 (2.526) |
| Promoter + 500bp | 0.039 | 0.169 (0.056) | 4.364 (1.442) |
| Repressed | 0.461 | 0.179 (0.145) | 0.388 (0.315) |
| Repressed + 500bp | 0.719 | 0.260 (0.078) | 0.362 (0.109) |
| Super Enhancer | 0.168 | 0.563 (0.061) | 3.341 (0.362) |
| Super Enhancer + 500bp | 0.172 | 0.615 (0.059) | 3.582 (0.346) |
| TFBS | 0.132 | 0.791 (0.221) | 5.970 (1.666) |
| TFBS + 500bp | 0.343 | 0.552 (0.182) | 1.607 (0.531) |
| Transcribed | 0.345 | 0.302 (0.144) | 0.875 (0.418) |
| Transcribed + 500bp | 0.763 | 0.596 (0.104) | 0.781 (0.136) |
| TSS | 0.018 | 0.341 (0.091) | 18.703 (4.996) |
| TSS + 500bp | 0.035 | 0.372 (0.080) | 10.679 (2.290) |
| 3-prime UTR | 0.011 | 0.073 (0.047) | 6.588 (4.219) |
| 3-prime UTR + 500bp | 0.027 | 0.107 (0.055) | 3.958 (2.056) |
| 5-prime UTR | 0.005 | 0.019 (0.044) | 3.562 (8.104) |
| 5-prime UTR + 500bp | 0.028 | 0.150 (0.063) | 5.381 (2.248) |
| Weak Enhancer | 0.021 | 0.230 (0.116) | 10.914 (5.510) |
| Weak Enhancer + 500bp | 0.089 | 0.424 (0.108) | 4.762 (1.214) |

**(S4.P).** Proportion of heritability and enrichment for different functional categories for Crohn’s disease.

(S4.O) Proportion of heritability and enrichment for different functional categories for Rheumatoid arthritis.
| Annotation | Prop. SNPs | Prop. $h^2$ | Enrichment |
| --- | --- | --- | --- |
| Coding | 0.015 | 0.125 (0.071) | 8.551 (4.851) |
| Coding + 500bp | 0.065 | 0.352 (0.065) | 5.453 (1.007) |
| Conserved | 0.026 | 0.069 (0.072) | 2.640 (2.766) |
| Conserved + 500bp | 0.333 | 0.919 (0.098) | 2.765 (0.296) |
| CTCF | 0.024 | -0.050 (0.087) | -2.081 (3.645) |
| CTCF + 500bp | 0.071 | -0.127 (0.083) | -1.781 (1.174) |
| DGF | 0.138 | 0.770 (0.182) | 5.600 (1.324) |
| DGF + 500bp | 0.542 | 1.088 (0.090) | 2.009 (0.166) |
| DHS peaks | 0.112 | 0.303 (0.136) | 2.713 (1.220) |
| DHS | 0.168 | 0.154 (0.177) | 0.915 (1.057) |
| DHS + 500bp | 0.499 | 0.713 (0.126) | 1.430 (0.253) |
| FANTOM5 Enhancer | 0.004 | 0.169 (0.042) | 39.042 (9.623) |
| FANTOM5 Enhancer + 500bp | 0.019 | 0.428 (0.073) | 22.468 (3.853) |
| Enhancer | 0.063 | 0.237 (0.096) | 3.740 (1.519) |
| Enhancer + 500bp | 0.154 | 0.554 (0.106) | 3.600 (0.691) |
| Fetal DHS | 0.085 | 0.398 (0.133) | 4.693 (1.570) |
| Fetal DHS + 500bp | 0.285 | 0.686 (0.131) | 2.406 (0.461) |
| H3K27ac<sup>19</sup> | 0.391 | 0.838 (0.049) | 2.143 (0.126) |
| H3K27ac<sup>19</sup> + 500bp | 0.423 | 0.840 (0.060) | 1.987 (0.141) |
| H3K27ac<sup>18</sup> | 0.269 | 0.768 (0.121) | 2.849 (0.450) |
| H3K27ac<sup>18</sup> + 500bp | 0.336 | 0.748 (0.074) | 2.227 (0.221) |
| H3K4me1 peaks | 0.171 | 0.506 (0.169) | 2.953 (0.987) |
| H3K4me1 | 0.427 | 0.742 (0.132) | 1.739 (0.310) |
| H3K4me1 + 500bp | 0.609 | 1.004 (0.068) | 1.648 (0.111) |
| H3K4me3 peaks | 0.042 | 0.318 (0.157) | 7.621 (3.765) |
| H3K4me3 | 0.133 | 0.675 (0.147) | 5.064 (1.100) |
| H3K4me3 + 500bp | 0.255 | 0.669 (0.126) | 2.618 (0.492) |
| H3K9ac peaks | 0.039 | 0.082 (0.113) | 2.109 (2.926) |
| H3K9ac | 0.126 | 0.231 (0.122) | 1.834 (0.964) |
| H3K9ac + 500bp | 0.231 | 0.565 (0.101) | 2.450 (0.438) |
| Intron | 0.387 | 0.400 (0.068) | 1.032 (0.176) |
| Intron + 500bp | 0.397 | 0.504 (0.054) | 1.268 (0.135) |
| PromoterFlanking | 0.008 | -0.054 (0.045) | -6.361 (5.315) |
| PromoterFlanking + 500bp | 0.033 | 0.018 (0.066) | 0.535 (1.959) |
| Promoter | 0.031 | 0.146 (0.065) | 4.685 (2.075) |
| Promoter + 500bp | 0.039 | 0.137 (0.047) | 3.553 (1.223) |
| Repressed | 0.461 | 0.102 (0.114) | 0.222 (0.248) |
| Repressed + 500bp | 0.719 | 0.392 (0.052) | 0.545 (0.072) |
| Super Enhancer | 0.168 | 0.554 (0.050) | 3.289 (0.297) |
| Super Enhancer + 500bp | 0.172 | 0.594 (0.046) | 3.462 (0.270) |
| TFBS | 0.132 | 0.614 (0.195) | 4.635 (1.474) |
| TFBS + 500bp | 0.343 | 0.730 (0.133) | 2.126 (0.386) |
| Transcribed | 0.345 | 0.402 (0.124) | 1.163 (0.359) |
| Transcribed + 500bp | 0.763 | 0.747 (0.081) | 0.978 (0.106) |
| TSS | 0.018 | 0.248 (0.073) | 13.613 (4.019) |
| TSS + 500bp | 0.035 | 0.398 (0.070) | 11.424 (2.013) |
| 3-prime UTR | 0.011 | 0.021 (0.035) | 1.909 (3.148) |
| 3-prime UTR + 500bp | 0.027 | 0.171 (0.076) | 6.347 (2.827) |
| 5-prime UTR | 0.005 | 0.040 (0.045) | 7.436 (8.298) |
| 5-prime UTR + 500bp | 0.028 | 0.151 (0.047) | 5.413 (1.703) |
| Weak Enhancer | 0.021 | 0.101 (0.069) | 4.797 (3.258) |
| Weak Enhancer + 500bp | 0.089 | 0.258 (0.084) | 2.903 (0.940) |

**(S4.Q).** Proportion of heritability and enrichment for different functional categories for Ulcerative colitis.

(S4.P) Proportion of heritability and enrichment for different functional categories for Crohn's disease.
| Annotation | Prop. SNPs | Prop. $h^2$ | Enrichment |
| --- | --- | --- | --- |
| Coding | 0.015 | 0.172 (0.088) | 11.727 (6.022) |
| Coding + 500bp | 0.065 | 0.421 (0.079) | 6.529 (1.218) |
| Conserved | 0.026 | 0.203 (0.091) | 7.786 (3.478) |
| Conserved + 500bp | 0.333 | 0.878 (0.123) | 2.639 (0.369) |
| CTCF | 0.024 | -0.101 (0.106) | -4.218 (4.430) |
| CTCF + 500bp | 0.071 | -0.077 (0.111) | -1.087 (1.559) |
| DGF | 0.138 | 0.901 (0.247) | 6.546 (1.795) |
| DGF + 500bp | 0.542 | 1.117 (0.134) | 2.063 (0.248) |
| DHS peaks | 0.112 | 0.706 (0.218) | 6.319 (1.950) |
| DHS | 0.168 | 0.416 (0.236) | 2.481 (1.409) |
| DHS + 500bp | 0.499 | 0.815 (0.165) | 1.635 (0.330) |
| FANTOM5 Enhancer | 0.004 | 0.181 (0.049) | 41.783 (11.190) |
| FANTOM5 Enhancer + 500bp | 0.019 | 0.285 (0.076) | 14.939 (3.999) |
| Enhancer | 0.063 | 0.348 (0.135) | 5.490 (2.133) |
| Enhancer + 500bp | 0.154 | 0.699 (0.138) | 4.544 (0.897) |
| Fetal DHS | 0.085 | 0.360 (0.186) | 4.243 (2.200) |
| Fetal DHS + 500bp | 0.285 | 0.550 (0.170) | 1.929 (0.596) |
| H3K27ac<sup>19</sup> | 0.391 | 0.989 (0.084) | 2.527 (0.214) |
| H3K27ac<sup>19</sup> + 500bp | 0.423 | 1.031 (0.088) | 2.440 (0.209) |
| H3K27ac<sup>18</sup> | 0.269 | 0.848 (0.163) | 3.148 (0.603) |
| H3K27ac<sup>18</sup> + 500bp | 0.336 | 0.813 (0.132) | 2.420 (0.394) |
| H3K4me1 peaks | 0.171 | 0.710 (0.243) | 4.142 (1.418) |
| H3K4me1 | 0.427 | 0.774 (0.173) | 1.814 (0.406) |
| H3K4me1 + 500bp | 0.609 | 0.911 (0.078) | 1.495 (0.128) |
| H3K4me3 peaks | 0.042 | 0.264 (0.174) | 6.318 (4.161) |
| H3K4me3 | 0.133 | 0.709 (0.179) | 5.321 (1.345) |
| H3K4me3 + 500bp | 0.255 | 0.707 (0.137) | 2.767 (0.536) |
| H3K9ac peaks | 0.039 | 0.276 (0.144) | 7.125 (3.713) |
| H3K9ac | 0.126 | 0.584 (0.172) | 4.634 (1.367) |
| H3K9ac + 500bp | 0.231 | 0.692 (0.122) | 3.002 (0.531) |
| Intron | 0.387 | 0.364 (0.082) | 0.941 (0.210) |
| Intron + 500bp | 0.397 | 0.506 (0.055) | 1.275 (0.138) |
| PromoterFlanking | 0.008 | 0.030 (0.055) | 3.586 (6.532) |
| PromoterFlanking + 500bp | 0.033 | 0.049 (0.080) | 1.464 (2.381) |
| Promoter | 0.031 | 0.150 (0.072) | 4.799 (2.296) |
| Promoter + 500bp | 0.039 | 0.185 (0.054) | 4.797 (1.392) |
| Repressed | 0.461 | 0.065 (0.160) | 0.140 (0.347) |
| Repressed + 500bp | 0.719 | 0.275 (0.072) | 0.383 (0.100) |
| Super Enhancer | 0.168 | 0.567 (0.062) | 3.365 (0.367) |
| Super Enhancer + 500bp | 0.172 | 0.616 (0.057) | 3.593 (0.331) |
| TFBS | 0.132 | 1.144 (0.283) | 8.637 (2.134) |
| TFBS + 500bp | 0.343 | 0.660 (0.173) | 1.921 (0.503) |
| Transcribed | 0.345 | 0.446 (0.162) | 1.291 (0.470) |
| Transcribed + 500bp | 0.763 | 0.677 (0.107) | 0.888 (0.140) |
| TSS | 0.018 | 0.209 (0.065) | 11.447 (3.569) |
| TSS + 500bp | 0.035 | 0.307 (0.076) | 8.827 (2.188) |
| 3-prime UTR | 0.011 | 0.034 (0.053) | 3.111 (4.792) |
| 3-prime UTR + 500bp | 0.027 | 0.176 (0.065) | 6.524 (2.425) |
| 5-prime UTR | 0.005 | 0.097 (0.043) | 17.831 (7.949) |
| 5-prime UTR + 500bp | 0.028 | 0.204 (0.073) | 7.325 (2.612) |
| Weak Enhancer | 0.021 | 0.057 (0.099) | 2.723 (4.693) |
| Weak Enhancer + 500bp | 0.089 | 0.328 (0.106) | 3.690 (1.190) |

**Table S5:** Proportion of heritability and enrichment for different functional categories, meta-analyzed over nine traits, including derived allele frequency bins in the model (Methods).

| Annotation | Prop. SNPs | Prop. $h^2$ | Enrichment |
| --- | --- | --- | --- |
| Coding | 0.015 | 0.108 (0.013) | 7.357 (0.859) |
| Coding + 500bp | 0.065 | 0.192 (0.030) | 2.979 (0.469) |
| Conserved | 0.026 | 0.359 (0.042) | 13.762 (1.603) |
| Conserved + 500bp | 0.333 | 0.656 (0.026) | 1.972 (0.080) |
| CTCF | 0.024 | -0.014 (0.019) | -0.578 (0.807) |
| CTCF + 500bp | 0.071 | 0.043 (0.020) | 0.601 (0.278) |
| DGF | 0.138 | 0.367 (0.097) | 2.669 (0.702) |
| DGF + 500bp | 0.542 | 0.747 (0.071) | 1.380 (0.132) |
| DHS peaks | 0.112 | 0.221 (0.065) | 1.978 (0.584) |
| DHS | 0.168 | 0.280 (0.071) | 1.666 (0.420) |
| DHS + 500bp | 0.499 | 0.756 (0.040) | 1.516 (0.081) |
| FANTOM5 Enhancer | 0.004 | -0.004 (0.009) | -0.853 (2.162) |
| FANTOM5 Enhancer + 500bp | 0.019 | 0.016 (0.018) | 0.815 (0.921) |
| Enhancer | 0.063 | 0.242 (0.051) | 3.825 (0.811) |
| Enhancer + 500bp | 0.154 | 0.349 (0.043) | 2.266 (0.280) |
| Fetal DHS | 0.085 | 0.236 (0.044) | 2.787 (0.517) |
| Fetal DHS + 500bp | 0.285 | 0.559 (0.059) | 1.960 (0.207) |
| H3K27ac<sup>19</sup> | 0.391 | 0.628 (0.055) | 1.605 (0.141) |
| H3K27ac<sup>19</sup> + 500bp | 0.423 | 0.664 (0.060) | 1.571 (0.142) |
| H3K27ac<sup>18</sup> | 0.269 | 0.485 (0.056) | 1.798 (0.207) |
| H3K27ac<sup>18</sup> + 500bp | 0.336 | 0.607 (0.042) | 1.806 (0.124) |
| H3K4me1 peaks | 0.171 | 0.464 (0.041) | 2.711 (0.241) |
| H3K4me1 | 0.427 | 0.807 (0.066) | 1.891 (0.156) |
| H3K4me1 + 500bp | 0.609 | 0.899 (0.041) | 1.476 (0.067) |
| H3K4me3 peaks | 0.042 | 0.166 (0.026) | 3.966 (0.616) |
| H3K4me3 | 0.133 | 0.345 (0.045) | 2.587 (0.335) |
| H3K4me3 + 500bp | 0.255 | 0.488 (0.058) | 1.909 (0.227) |
| H3K9ac peaks | 0.039 | 0.259 (0.025) | 6.674 (0.635) |
| H3K9ac | 0.126 | 0.409 (0.057) | 3.240 (0.452) |
| H3K9ac + 500bp | 0.231 | 0.502 (0.041) | 2.178 (0.178) |
| Intron | 0.387 | 0.467 (0.014) | 1.206 (0.035) |
| Intron + 500bp | 0.397 | 0.528 (0.015) | 1.329 (0.037) |
| PromoterFlanking | 0.008 | 0.010 (0.011) | 1.137 (1.346) |
| PromoterFlanking + 500bp | 0.033 | 0.083 (0.018) | 2.488 (0.543) |
| Promoter | 0.031 | 0.091 (0.016) | 2.923 (0.521) |
| Promoter + 500bp | 0.039 | 0.079 (0.017) | 2.045 (0.441) |
| Repressed | 0.461 | 0.268 (0.065) | 0.581 (0.141) |
| Repressed + 500bp | 0.719 | 0.443 (0.049) | 0.616 (0.068) |
| Super Enhancer | 0.168 | 0.303 (0.036) | 1.797 (0.214) |
| Super Enhancer + 500bp | 0.172 | 0.316 (0.038) | 1.843 (0.221) |
| TFBS | 0.132 | 0.440 (0.065) | 3.321 (0.492) |
| TFBS + 500bp | 0.343 | 0.481 (0.054) | 1.399 (0.156) |
| Transcribed | 0.345 | 0.430 (0.039) | 1.244 (0.112) |
| Transcribed + 500bp | 0.763 | 0.726 (0.028) | 0.951 (0.037) |
| TSS | 0.018 | 0.105 (0.024) | 5.768 (1.303) |
| TSS + 500bp | 0.035 | 0.171 (0.029) | 4.907 (0.839) |
| 3-prime UTR | 0.011 | 0.053 (0.010) | 4.781 (0.872) |
| 3-prime UTR + 500bp | 0.027 | 0.075 (0.011) | 2.792 (0.422) |
| 5-prime UTR | 0.005 | 0.029 (0.008) | 5.309 (1.402) |
| 5-prime UTR + 500bp | 0.028 | 0.067 (0.011) | 2.395 (0.384) |
| Weak Enhancer | 0.021 | 0.070 (0.024) | 3.326 (1.141) |
| Weak Enhancer + 500bp | 0.089 | 0.190 (0.031) | 2.138 (0.343) |

**Table S6:**
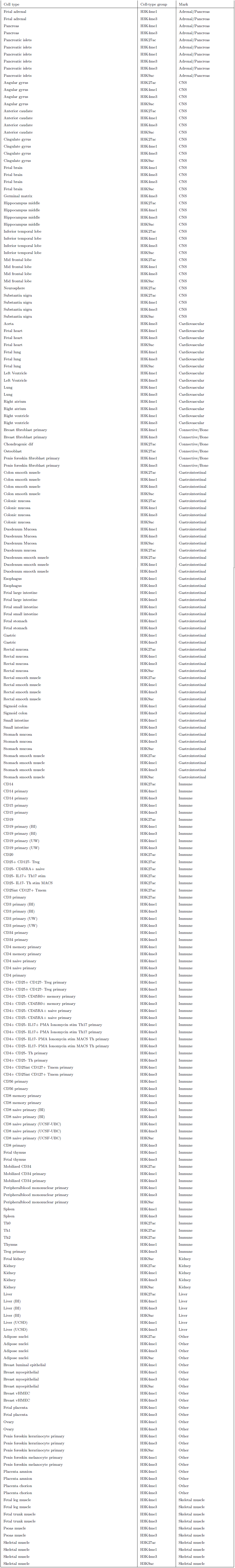
Cell types used in the cell-type-specific analysis. When the same cell type in the same histone mark from more than one institution was used, the institution is given in parentheses.

**(S7.A).** Enrichment of top cell types for Height. * = significant at FDR < 0.05. ** = significant at p < 0.05 after correcting for multiple hypotheses.

| Cell type | Cell type group | Mark | $-\log_{10}(p)$ |
| --- | --- | --- | --- |
| Chondrogenic dif** | Connective/Bone | H3K27ac | 6.81 |
| Penis foreskin fibroblast primary** | Connective/Bone | H3K4me1 | 6.43 |
| Fetal lung** | Cardiovascular | H3K4me1 | 6.34 |
| Fetal stomach** | GI | H3K4me1 | 5.48 |
| Colon smooth muscle* | GI | H3K4me1 | 4.64 |
| Aorta* | Cardiovascular | H3K4me3 | 4.64 |
| Fetal lung* | Cardiovascular | H3K9ac | 4.31 |
| Stomach smooth muscle* | GI | H3K4me3 | 4.26 |
| Osteoblast* | Connective/Bone | H3K27ac | 4.04 |
| Penis foreskin fibroblast primary* | Connective/Bone | H3K4me3 | 3.96 |
| Stomach smooth muscle* | GI | H3K4me1 | 3.94 |
| Fetal leg muscle* | Skeletal Muscle | H3K4me3 | 3.91 |
| Fetal trunk muscle* | Skeletal Muscle | H3K4me3 | 3.72 |
| Rectal smooth muscle* | GI | H3K4me3 | 3.57 |
| Fetal lung* | Cardiovascular | H3K4me3 | 3.37 |
| Rectal smooth muscle* | GI | H3K4me1 | 3.32 |
| Fetal placenta* | Other | H3K4me3 | 3.26 |
| Adipose nuclei* | Other | H3K4me1 | 3.1 |
| Ovary* | Other | H3K4me1 | 3.06 |
| Fetal large intestine* | GI | H3K4me3 | 3.05 |
| Placenta chorion* | Other | H3K4me3 | 2.98 |
| CD34 primary* | Immune | H3K4me1 | 2.96 |
| Penis foreskin melanocyte primary* | Other | H3K4me1 | 2.95 |
| Skeletal muscle* | Skeletal Muscle | H3K9ac | 2.93 |
| Mobilized CD34 primary* | Immune | H3K4me1 | 2.88 |
| Fetal stomach* | GI | H3K4me3 | 2.87 |
| Mobilized CD34 primary* | Immune | H3K4me3 | 2.87 |
| Fetal adrenal* | Adrenal/Pancreas | H3K4me3 | 2.86 |
| Breast fibroblast primary* | Connective/Bone | H3K4me3 | 2.85 |
| Duodenum smooth muscle* | GI | H3K4me1 | 2.81 |
| Colon smooth muscle* | GI | H3K4me3 | 2.76 |
| Ovary* | Other | H3K4me3 | 2.7 |
| Fetal brain* | CNS | H3K4me3 | 2.62 |
| Skeletal muscle* | Skeletal Muscle | H3K4me1 | 2.6 |
| Fetal small intestine* | GI | H3K4me3 | 2.6 |
| Colon smooth muscle* | GI | H3K27ac | 2.57 |
| Lung* | Cardiovascular | H3K4me3 | 2.56 |
| Liver (UCSD)* | Liver | H3K4me3 | 2.53 |
| Esophagus* | GI | H3K4me3 | 2.53 |
| Placenta amnion* | Other | H3K4me3 | 2.48 |
| Right ventricle* | Cardiovascular | H3K4me3 | 2.47 |
| Sigmoid colon* | GI | H3K4me3 | 2.44 |
| Fetal leg muscle* | Skeletal Muscle | H3K4me1 | 2.44 |
| Colonic mucosa* | GI | H3K4me3 | 2.18 |
| Right atrium* | Cardiovascular | H3K4me3 | 2.14 |
| CD34 primary* | Immune | H3K4me3 | 2.13 |
| Gastric* | GI | H3K4me3 | 2.08 |
| Skeletal muscle* | Skeletal Muscle | H3K4me3 | 2.07 |
| Lung* | Cardiovascular | H3K4me1 | 2.06 |
| Pancreatic islets* | Adrenal/Pancreas | H3K4me3 | 2.05 |
| Adipose nuclei* | Other | H3K9ac | 2.01 |
| Right atrium* | Cardiovascular | H3K4me1 | 1.98 |
| Stomach smooth muscle* | GI | H3K27ac | 1.96 |
| Rectal smooth muscle* | GI | H3K27ac | 1.95 |
| Breast fibroblast primary* | Connective/Bone | H3K4me1 | 1.93 |
| Germinal matrix* | CNS | H3K4me3 | 1.92 |
| Small intestine* | GI | H3K4me3 | 1.91 |
| Fetal placenta* | Other | H3K4me1 | 1.91 |

**(S7.B).** Enrichment of top cell types for BMI. * = significant at FDR < 0.05. ** = significant at p < 0.05 after correcting for multiple hypotheses.

| Cell type | Cell type group | Mark | $-\log_{10}(p)$ |
| --- | --- | --- | --- |
| Fetal brain* | CNS | H3K4me3 | 4.48 |
| Penis foreskin fibroblast primary* | Connective/Bone | H3K4me3 | 4.43 |
| Inferior temporal lobe* | CNS | H3K4me1 | 4.3 |
| Mid frontal lobe* | CNS | H3K9ac | 4.25 |
| Anterior caudate* | CNS | H3K4me3 | 4.25 |
| Mid frontal lobe* | CNS | H3K27ac | 3.96 |
| Anterior caudate* | CNS | H3K9ac | 3.91 |
| Cingulate gyrus* | CNS | H3K4me1 | 3.73 |
| Inferior temporal lobe* | CNS | H3K4me3 | 3.73 |
| Penis foreskin keratinocyte primary* | Other | H3K9ac | 3.72 |
| Mid frontal lobe* | CNS | H3K4me3 | 3.71 |
| Hippocampus middle* | CNS | H3K4me1 | 3.66 |
| Inferior temporal lobe* | CNS | H3K9ac | 3.59 |
| Fetal brain* | CNS | H3K9ac | 3.57 |
| Hippocampus middle* | CNS | H3K9ac | 3.47 |
| Cingulate gyrus* | CNS | H3K9ac | 3.46 |
| Hippocampus middle* | CNS | H3K4me3 | 3.4 |
| Germinal matrix* | CNS | H3K4me3 | 3.4 |
| Cingulate gyrus* | CNS | H3K4me3 | 3.4 |
| Anterior caudate* | CNS | H3K4me1 | 3.31 |
| Substantia nigra* | CNS | H3K4me3 | 3.24 |
| Angular gyrus* | CNS | H3K27ac | 3.05 |
| Penis foreskin melanocyte primary* | Other | H3K4me3 | 2.83 |
| Angular gyrus* | CNS | H3K4me3 | 2.76 |
| Substantia nigra* | CNS | H3K4me1 | 2.75 |
| Pancreatic islets* | Adrenal/Pancreas | H3K4me3 | 2.6 |
| Cingulate gyrus* | CNS | H3K27ac | 2.57 |
| Fetal adrenal* | Adrenal/Pancreas | H3K4me3 | 2.57 |
| Angular gyrus* | CNS | H3K9ac | 2.51 |
| Inferior temporal lobe* | CNS | H3K27ac | 2.39 |
| Breast myoepithelial* | Other | H3K4me3 | 2.35 |
| Substantia nigra* | CNS | H3K9ac | 2.26 |
| Substantia nigra* | CNS | H3K27ac | 2.22 |
| Hippocampus middle* | CNS | H3K27ac | 2.07 |

**(S7.C).**
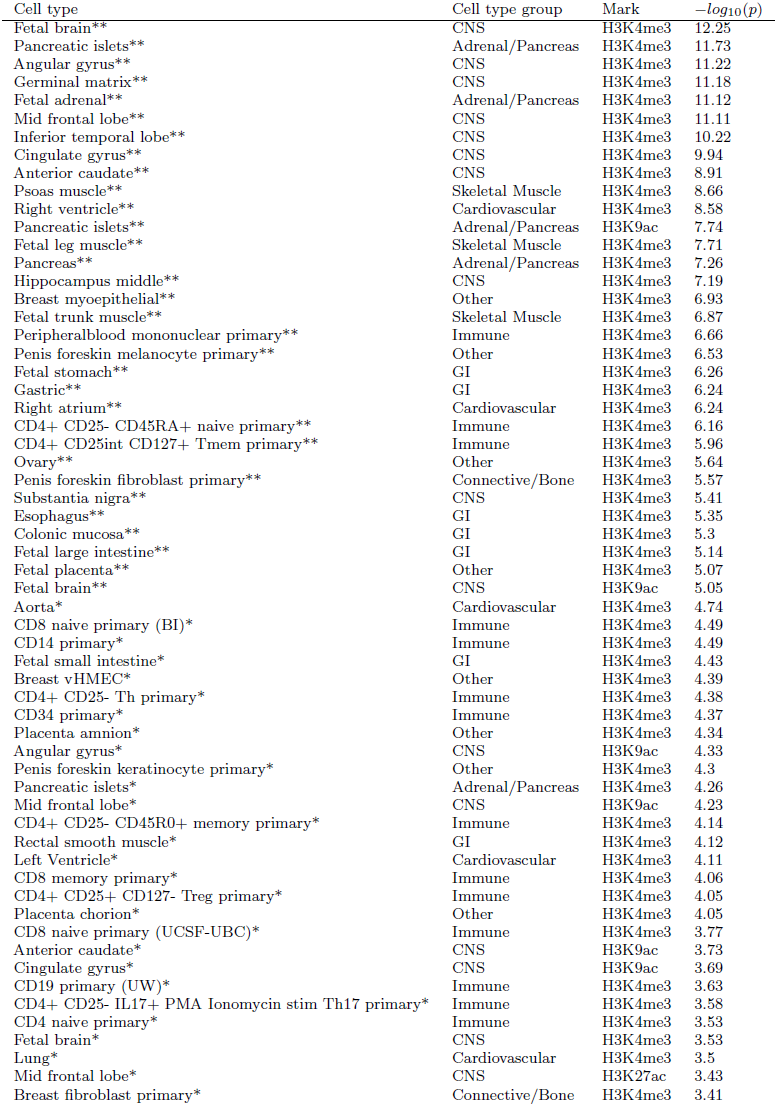
Enrichment of top cell types for Age at menarche. * = significant at FDR < 0.05. ** = significant at p < 0.05 after correcting for multiple hypotheses.

**(S7.D).** Enrichment of top cell types for LDL. * = significant at FDR < 0.05. ** = significant at p < 0.05 after correcting for multiple hypotheses.

| Cell type | Cell type group | Mark | $-\log_{10}(p)$ |
| --- | --- | --- | --- |
| Liver (BI)* | Liver | H3K4me1 | 4.76 |
| Fetal adrenal* | Adrenal/Pancreas | H3K4me1 | 3.41 |
| CD14 primary* | Immune | H3K4me1 | 3.33 |
| Liver* | Liver | H3K27ac | 2.97 |
| Adipose nuclei | Other | H3K9ac | 2.71 |

**(S7.E).** Enrichment of top cell types for HDL. * = significant at FDR < 0.05. ** = signidicant at p < 0.05 after correcting for multiple hypotheses.

| Cell type | Cell type group | Mark | $-\log_{10}(p)$ |
| --- | --- | --- | --- |
| Liver (BI)* | Liver | H3K4me1 | 4.51 |
| Adipose nuclei* | Other | H3K4me1 | 4.26 |
| Liver* | Liver | H3K27ac | 3.61 |
| Adipose nuclei* | Other | H3K9ac | 3.34 |
| Adipose nuclei* | Other | H3K4me3 | 3.08 |
| CD14 primary* | Immune | H3K4me1 | 2.86 |
| Adipose nuclei* | Other | H3K27ac | 2.84 |
| Liver (BI)* | Liver | H3K9ac | 2.74 |
| Liver (BI)* | Liver | H3K4me3 | 2.66 |

**(S7.F).** Enrichment of top cell types for Triglycerides. * = significant at FDR < 0.05. ** = significant at p < 0.05 after correcting for multiple hypotheses.

| Cell type | Cell type group | Mark | $-\log_{10}(p)$ |
| --- | --- | --- | --- |
| Liver (BI)* | Liver | H3K4me1 | 3.99 |
| Liver* | Liver | H3K27ac | 3.66 |
| Liver (BI)* | Liver | H3K9ac | 3.02 |
| Duodenum Mucosa | GI | H3K4me3 | 2.71 |
| Liver (UCSD) | Liver | H3K4me3 | 2.68 |

**(S7.G).**
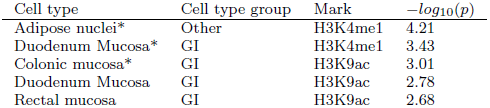
Enrichment of top cell types for Coronary artery disease. * = significant at FDR < 0.05. ** = significant at p < 0.05 after correcting for multiple hypotheses.

**(S7.H).** Enrichment of top cell types for Type 2 Diabetes. * = significant at FDR < 0.05. ** = significant at p < 0.05 after correcting for multiple hypotheses.

| Cell type | Cell type group | Mark | $-\log_{10}(p)$ |
| --- | --- | --- | --- |
| Pancreatic islets | Adrenal/Pancreas | H3K4me3 | 2.87 |
| Pancreatic islets | Adrenal/Pancreas | H3K27ac | 2.73 |
| Fetal large intestine | GI | H3K4me1 | 2.49 |
| Fetal small intestine | GI | H3K4me1 | 2.31 |
| Adipose nuclei | Other | H3K9ac | 2.27 |

**(S7.I).** Enrichment of top cell types for Fasting Glucose. * = significant at FDR < 0.05. ** = significant at p < 0.05 after correcting for multiple hypotheses.

| Cell type | Cell type group | Mark | $-\log_{10}(p)$ |
| --- | --- | --- | --- |
| Pancreatic islets* | Adrenal/Pancreas | H3K27ac | 3.93 |
| Pancreatic islets* | Adrenal/Pancreas | H3K4me1 | 3.1 |
| Pancreatic islets | Adrenal/Pancreas | H3K4me3 | 2.93 |
| Pancreatic islets | Adrenal/Pancreas | H3K4me3 | 2.25 |
| Fetal small intestine | GI | H3K4me1 | 2.18 |

**(S7.J).** Enrichment of top cell types for Schizophrenia. * = significant at FDR < 0.05. ** = significant at p < 0.05 after correcting for multiple hypotheses.

| Cell type | Cell type group | Mark | $-\log_{10}(p)$ |
| --- | --- | --- | --- |
| Fetal brain** | CNS | H3K4me3 | 18.51 |
| Mid frontal lobe** | CNS | H3K4me3 | 14.44 |
| Germinal matrix** | CNS | H3K4me3 | 12.68 |
| Mid frontal lobe** | CNS | H3K9ac | 11.27 |
| Angular gyrus** | CNS | H3K4me3 | 10.89 |
| Inferior temporal lobe** | CNS | H3K4me3 | 10.77 |
| Cingulate gyrus** | CNS | H3K9ac | 10.27 |
| Fetal brain** | CNS | H3K9ac | 10.24 |
| Anterior caudate** | CNS | H3K4me3 | 9.66 |
| Cingulate gyrus** | CNS | H3K4me3 | 9.34 |
| Pancreatic islets** | Adrenal/Pancreas | H3K4me3 | 8.65 |
| Anterior caudate** | CNS | H3K9ac | 8.5 |
| Angular gyrus** | CNS | H3K9ac | 8.33 |
| Mid frontal lobe** | CNS | H3K27ac | 8.1 |
| Anterior caudate** | CNS | H3K4me1 | 7.92 |
| Inferior temporal lobe** | CNS | H3K4me1 | 7.43 |
| Psoas muscle** | Skeletal Muscle | H3K4me3 | 7.38 |
| Fetal brain** | CNS | H3K4me1 | 7.21 |
| Inferior temporal lobe** | CNS | H3K9ac | 7.03 |
| Hippocampus middle** | CNS | H3K9ac | 6.03 |
| Pancreatic islets** | Adrenal/Pancreas | H3K9ac | 5.79 |
| Penis foreskin melanocyte primary** | Other | H3K4me3 | 5.68 |
| Angular gyrus** | CNS | H3K27ac | 5.63 |
| Cingulate gyrus** | CNS | H3K4me1 | 5.55 |
| Hippocampus middle** | CNS | H3K4me3 | 5.55 |
| CD34 primary** | Immune | H3K4me3 | 5.33 |
| Sigmoid colon** | GI | H3K4me3 | 5.3 |
| Fetal adrenal** | Adrenal/Pancreas | H3K4me3 | 5.2 |
| Inferior temporal lobe** | CNS | H3K27ac | 5.08 |
| Peripheral blood mononuclear primary** | Immune | H3K4me3 | 5.03 |
| Gastric** | GI | H3K4me3 | 4.93 |
| Substantia nigra* | CNS | H3K4me3 | 4.71 |
| Fetal brain* | CNS | H3K4me3 | 4.58 |
| Hippocampus middle* | CNS | H3K4me1 | 4.48 |
| Ovary* | Other | H3K4me3 | 4.19 |
| CD19 primary (UW)* | Immune | H3K4me3 | 4.15 |
| Small intestine* | GI | H3K4me3 | 4.07 |
| Lung* | Cardiovascular | H3K4me3 | 3.93 |
| Fetal stomach* | GI | H3K4me3 | 3.89 |
| Fetal leg muscle* | Skeletal Muscle | H3K4me3 | 3.82 |
| Spleen* | Immune | H3K4me3 | 3.77 |
| Breast fibroblast primary* | Connective/Bone | H3K4me3 | 3.69 |
| Right ventricle* | Cardiovascular | H3K4me3 | 3.67 |
| CD4+ CD25- Th primary* | Immune | H3K4me3 | 3.66 |
| CD4+ CD25- IL17- PMA Ionomycin stim MACS Th primary* | Immune | H3K4me1 | 3.66 |
| CD8 naive primary (UCSF-UBC)* | Immune | H3K4me3 | 3.65 |
| Pancreas* | Adrenal/Pancreas | H3K4me3 | 3.63 |
| CD4+ CD25- Th primary* | Immune | H3K4me1 | 3.56 |
| CD4+ CD25- CD45RA+ naive primary* | Immune | H3K4me1 | 3.56 |
| Colonic mucosa* | GI | H3K4me3 | 3.49 |
| Right atrium* | Cardiovascular | H3K4me3 | 3.48 |
| Fetal trunk muscle* | Skeletal Muscle | H3K4me3 | 3.47 |
| CD4+ CD25int CD127+ Tmem primary* | Immune | H3K4me3 | 3.46 |
| Substantia nigra* | CNS | H3K9ac | 3.44 |
| Placenta amnion* | Other | H3K4me3 | 3.38 |
| Breast myoepithelial* | Other | H3K9ac | 3.26 |
| CD8 naive primary (BI)* | Immune | H3K4me1 | 3.24 |
| Substantia nigra* | CNS | H3K4me1 | 3.18 |
| Cingulate gyrus* | CNS | H3K27ac | 3.1 |
| CD4+ CD25- CD45RA+ naive primary* | Immune | H3K4me3 | 3.06 |

**(S7.K).**
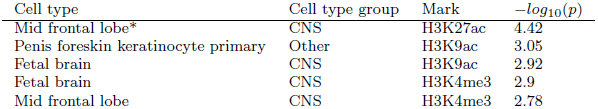
Enrichment of top cell types for Bipolar disorder. * = significant at FDR < 0.05. ** = significant at p < 0.05 after correcting for multiple hypotheses.

**(S7.L).** Enrichment of top cell types for Anorexia. * = significant at FDR < 0.05. ** = significant at p < 0.05 after correcting for multiple hypotheses.

| Cell type | Cell type group | Mark | $-\log_{10}(p)$ |
| --- | --- | --- | --- |
| Angular gyrus | CNS | H3K9ac | 2.61 |
| Mid frontal lobe | CNS | H3K9ac | 2.38 |
| Mid frontal lobe | CNS | H3K4me1 | 2.36 |
| Anterior caudate | CNS | H3K9ac | 2.28 |
| Cingulate gyrus | CNS | H3K9ac | 2.22 |

**(S7.M).** Enrichment of top cell types for Years of education. * = significant at FDR < 0.05. ** = significant at p < 0.05 after correcting for multiple hypotheses.

| Cell type | Cell type group | Mark | $-\log_{10}(p)$ |
| --- | --- | --- | --- |
| Angular gyrus** | CNS | H3K4me3 | 6.63 |
| Fetal brain** | CNS | H3K4me3 | 6.05 |
| Mid frontal lobe** | CNS | H3K4me3 | 5.99 |
| Anterior caudate** | CNS | H3K4me3 | 5.73 |
| Inferior temporal lobe** | CNS | H3K4me3 | 5.63 |
| CD56 primary** | Immune | H3K4me3 | 5.32 |
| Germinal matrix** | CNS | H3K4me3 | 5.29 |
| Mid frontal lobe** | CNS | H3K9ac | 5.26 |
| Cingulate gyrus** | CNS | H3K9ac | 4.98 |
| Cingulate gyrus** | CNS | H3K4me3 | 4.94 |
| CD8 naive primary (UCSF-UBC)* | Immune | H3K4me3 | 4.88 |
| Penis foreskin melanocyte primary* | Other | H3K4me3 | 4.73 |
| Mid frontal lobe* | CNS | H3K27ac | 4.43 |
| Peripheralblood mononuclear primary* | Immune | H3K4me3 | 4.39 |
| CD34 primary* | Immune | H3K4me3 | 4.12 |
| Fetal brain* | CNS | H3K9ac | 4.03 |
| CD14 primary* | Immune | H3K4me3 | 4.01 |
| Inferior temporal lobe* | CNS | H3K9ac | 3.98 |
| Angular gyrus* | CNS | H3K9ac | 3.96 |
| Sigmoid colon* | GI | H3K4me3 | 3.83 |
| Pancreatic islets* | Adrenal/Pancreas | H3K9ac | 3.83 |
| CD4+ CD25int CD127+ Tmem primary* | Immune | H3K4me3 | 3.72 |
| Anterior caudate* | CNS | H3K9ac | 3.65 |
| Hippocampus middle* | CNS | H3K4me3 | 3.65 |
| CD19 primary (UW)* | Immune | H3K4me3 | 3.63 |
| Small intestine* | GI | H3K4me3 | 3.52 |
| CD4+ CD25- Th primary* | Immune | H3K4me3 | 3.45 |
| Lung* | Cardiovascular | H3K4me3 | 3.19 |
| CD4+ CD25- CD45R0+ memory primary* | Immune | H3K4me3 | 3.19 |
| Hippocampus middle* | CNS | H3K9ac | 3.17 |
| Liver (UCSD)* | Liver | H3K4me3 | 3.08 |
| Fetal placenta* | Other | H3K4me3 | 3.08 |
| Fetal adrenal* | Adrenal/Pancreas | H3K4me3 | 3.04 |
| Right atrium* | Cardiovascular | H3K4me3 | 2.99 |
| Pancreatic islets* | Adrenal/Pancreas | H3K4me3 | 2.96 |
| CD8 naive primary (UCSF-UBC)* | Immune | H3K9ac | 2.95 |
| CD3 primary (BI)* | Immune | H3K4me3 | 2.94 |
| Angular gyrus* | CNS | H3K27ac | 2.93 |
| CD4+ CD25- CD45RA+ naive primary* | Immune | H3K4me3 | 2.93 |
| Gastric* | GI | H3K4me3 | 2.92 |
| CD4 naive primary* | Immune | H3K4me3 | 2.89 |
| CD8 memory primary* | Immune | H3K4me3 | 2.78 |
| CD3 primary (UW)* | Immune | H3K4me3 | 2.76 |
| Rectal smooth muscle* | GI | H3K4me3 | 2.73 |
| Fetal brain* | CNS | H3K4me3 | 2.67 |
| Esophagus* | GI | H3K4me3 | 2.66 |
| CD8 naive primary (BI)* | Immune | H3K4me3 | 2.58 |
| Left Ventricle* | Cardiovascular | H3K4me3 | 2.56 |
| CD19 primary (BI)* | Immune | H3K4me3 | 2.56 |
| Fetal thymus* | Immune | H3K4me3 | 2.52 |
| Breast vHMEC* | Other | H3K4me3 | 2.51 |
| CD8 primary* | Immune | H3K4me3 | 2.51 |
| Psoas muscle* | Skeletal Muscle | H3K4me3 | 2.51 |
| Peripheralblood mononuclear primary* | Immune | H3K9ac | 2.5 |
| Ovary* | Other | H3K4me3 | 2.47 |
| Pancreas* | Adrenal/Pancreas | H3K4me3 | 2.46 |
| Breast fibroblast primary* | Connective/Bone | H3K4me3 | 2.45 |
| CD4+ CD25+ CD127- Treg primary* | Immune | H3K4me3 | 2.36 |
| Placenta amnion* | Other | H3K4me3 | 2.34 |
| Right ventricle* | Cardiovascular | H3K4me3 | 2.33 |

**(S7.N).** Enrichment of top cell types for Ever smoked. * = significant at FDR < 0.05. ** = signigicant at p < 0.05 after correcting for multiple hypotheses.

| Cell type | Cell type group | Mark | $-\log_{10}(p)$ |
| --- | --- | --- | --- |
| Inferior temporal lobe* | CNS | H3K4me3 | 3.21 |
| Cingulate gyrus* | CNS | H3K27ac | 3.2 |
| Substantia nigra* | CNS | H3K27ac | 3.16 |
| Hippocampus middle* | CNS | H3K27ac | 3.13 |
| Breast myoepithelial* | Other | H3K9ac | 3.06 |
| Inferior temporal lobe* | CNS | H3K4me1 | 2.93 |
| Anterior caudate* | CNS | H3K27ac | 2.81 |
| Inferior temporal lobe* | CNS | H3K27ac | 2.81 |
| Angular gyrus* | CNS | H3K27ac | 2.77 |
| Pancreatic islets* | Adrenal/Pancreas | H3K4me1 | 2.55 |

**(S7.O).** Enrichment of top cell types for Rheumatoid arthritis. * = significant at FDR < 0.05. ** = significant at p < 0.05 after correcting for multiple hypotheses.

| Cell type | Cell type group | Mark | $-\log_{10}(p)$ |
| --- | --- | --- | --- |
| CD4+ CD25- IL17+ PMA Ionomycin stim Th17 primary** | Immune | H3K4me1 | 6.76 |
| CD4+ CD25- IL17- PMA Ionomycin stim MACS Th primary** | Immune | H3K4me1 | 6.11 |
| CD4+ CD25- CD45R0+ memory primary** | Immune | H3K4me1 | 5.92 |
| CD4 memory primary** | Immune | H3K4me1 | 5.88 |
| CD4+ CD25+ CD127- Treg primary** | Immune | H3K4me1 | 5.83 |
| CD25- IL17- Th stim MACS** | Immune | H3K27ac | 5.7 |
| Th2** | Immune | H3K27ac | 5.5 |
| CD8 memory primary** | Immune | H3K4me1 | 5.38 |
| CD4 naive primary** | Immune | H3K4me1 | 5.26 |
| CD4+ CD25- Th primary** | Immune | H3K4me1 | 5.25 |
| CD19 primary (UW)** | Immune | H3K4me1 | 5.25 |
| CD4+ CD25int CD127+ Tmem primary* | Immune | H3K4me1 | 4.88 |
| CD4+ CD25- CD45RA+ naive primary* | Immune | H3K4me1 | 4.75 |
| CD3 primary (BI)* | Immune | H3K4me1 | 4.64 |
| CD3 primary (UW)* | Immune | H3K4me1 | 4.63 |
| CD25- IL17+ Th17 stim* | Immune | H3K27ac | 4.55 |
| CD8 naive primary (UCSF-UBC)* | Immune | H3K4me1 | 4.49 |
| CD8 naive primary (BI)* | Immune | H3K4me1 | 4.45 |
| Th0* | Immune | H3K27ac | 4.09 |
| CD25+ CD127- Treg* | Immune | H3K27ac | 4.09 |
| Th1* | Immune | H3K27ac | 3.96 |
| CD19 primary (BI)* | Immune | H3K4me1 | 3.91 |
| CD56 primary* | Immune | H3K4me1 | 3.77 |
| Treg primary* | Immune | H3K4me3 | 3.63 |
| CD3 primary* | Immune | H3K27ac | 3.62 |
| CD20* | Immune | H3K27ac | 3.45 |
| CD4+ CD25- IL17- PMA Ionomycin stim MACS Th primary* | Immune | H3K4me3 | 3.45 |
| CD4+ CD25- IL17+ PMA Ionomycin stim Th17 primary* | Immune | H3K4me3 | 3.17 |
| CD4+ CD25+ CD127- Treg primary* | Immune | H3K4me3 | 3.1 |
| CD4+ CD25int CD127+ Tmem primary* | Immune | H3K4me3 | 2.76 |
| Peripheralblood mononuclear primary* | Immune | H3K9ac | 2.58 |
| CD25int CD127+ Tmem* | Immune | H3K27ac | 2.27 |
| CD4+ CD25- CD45R0+ memory primary* | Immune | H3K4me3 | 2.24 |
| CD4+ CD25- CD45RA+ naive primary* | Immune | H3K4me3 | 2.2 |
| CD8 memory primary* | Immune | H3K4me3 | 2.17 |
| CD19* | Immune | H3K27ac | 2.13 |
| CD4 memory primary* | Immune | H3K4me3 | 2.12 |
| Peripheralblood mononuclear primary* | Immune | H3K4me1 | 2.12 |
| CD4+ CD25- Th primary* | Immune | H3K4me3 | 1.98 |

**(S7.P).** Enrichment of top cell types for Crohn’s disease. * = significant at FDR < 0.05. ** = significant at p < 0.05 after correcting for multiple hypotheses.

| Cell type | Cell type group | Mark | $-\log_{10}(p)$ |
| --- | --- | --- | --- |
| CD4+ CD25- IL17+ PMA Ionomycin stim Th17 primary** | Immune | H3K4me1 | 7.59 |
| Th1** | Immune | H3K27ac | 6.54 |
| CD25- IL17+ Th17 stim** | Immune | H3K27ac | 6.5 |
| CD4+ CD25- IL17- PMA Ionomycin stim MACS Th primary** | Immune | H3K4me1 | 6.24 |
| CD4 memory primary** | Immune | H3K4me1 | 5.88 |
| Th2** | Immune | H3K27ac | 5.87 |
| CD4+ CD25- Th primary** | Immune | H3K4me1 | 5.59 |
| CD8 memory primary** | Immune | H3K4me1 | 5.13 |
| CD14 primary** | Immune | H3K4me1 | 5.03 |
| CD3 primary (UW)** | Immune | H3K4me1 | 4.96 |
| Th0* | Immune | H3K27ac | 4.8 |
| CD56 primary* | Immune | H3K4me1 | 4.8 |
| CD25- IL17- Th stim MACS* | Immune | H3K27ac | 4.72 |
| CD4+ CD25- CD45R0+ memory primary* | Immune | H3K4me1 | 4.7 |
| CD4 naive primary* | Immune | H3K4me1 | 4.51 |
| CD4+ CD25int CD127+ Tmem primary* | Immune | H3K4me1 | 4.44 |
| CD4+ CD25- CD45RA+ naive primary* | Immune | H3K4me1 | 4.36 |
| CD8 naive primary (BI)* | Immune | H3K4me1 | 4.31 |
| CD19 primary (UW)* | Immune | H3K4me1 | 4.26 |
| CD8 naive primary (UCSF-UBC)* | Immune | H3K4me1 | 4.2 |
| CD4+ CD25- IL17+ PMA Ionomycin stim Th17 primary* | Immune | H3K4me3 | 4.18 |
| CD19 primary (BI)* | Immune | H3K4me1 | 4.17 |
| CD3 primary (BI)* | Immune | H3K4me1 | 3.73 |
| CD4+ CD25+ CD127- Treg primary* | Immune | H3K4me1 | 3.62 |
| CD3 primary* | Immune | H3K27ac | 3.25 |
| CD4+ CD25- IL17- PMA Ionomycin stim MACS Th primary* | Immune | H3K4me3 | 3.16 |
| CD34 primary* | Immune | H3K4me1 | 2.87 |
| Peripheralblood mononuclear primary* | Immune | H3K9ac | 2.87 |
| CD15 primary* | Immune | H3K4me1 | 2.85 |
| Spleen* | Immune | H3K4me1 | 2.7 |
| CD4 primary* | Immune | H3K4me3 | 2.49 |
| Peripheralblood mononuclear primary* | Immune | H3K4me1 | 2.46 |
| CD8 primary* | Immune | H3K4me3 | 2.44 |
| CD14* | Immune | H3K27ac | 2.18 |
| CD4 memory primary* | Immune | H3K4me3 | 2.12 |
| Colonic mucosa* | GI | H3K4me1 | 2.11 |
| CD19 primary (BI)* | Immune | H3K4me3 | 2.1 |
| CD4 naive primary* | Immune | H3K4me3 | 2.09 |
| Mobilized CD34 primary* | Immune | H3K4me1 | 2.08 |
| CD25int CD127+ Tmem* | Immune | H3K27ac | 2.04 |

**(S7.Q).** Enrichment of top cell types for Ulcerative colitis. * = significant at FDR < 0.05. ** = significant at p < 0.05 after correcting for multiple hypotheses.

| Cell type | Cell type group | Mark | $-\log_{10}(p)$ |
| --- | --- | --- | --- |
| CD4+ CD25- IL17+ PMA Ionomycin stim Th17 primary** | Immune | H3K4me1 | 6.37 |
| CD25- IL17+ Th17 stim** | Immune | H3K27ac | 5.53 |
| CD4+ CD25- CD45R0+ memory primary** | Immune | H3K4me1 | 5.48 |
| CD4 memory primary** | Immune | H3K4me1 | 5.19 |
| CD4+ CD25+ CD127- Treg primary* | Immune | H3K4me1 | 4.88 |
| CD4+ CD25- IL17- PMA Ionomycin stim MACS Th primary* | Immune | H3K4me1 | 4.73 |
| CD3 primary* | Immune | H3K27ac | 4.72 |
| Th2* | Immune | H3K27ac | 4.5 |
| CD25+ CD127- Treg* | Immune | H3K27ac | 4.42 |
| Colonic mucosa* | GI | H3K4me1 | 4.17 |
| Spleen* | Immune | H3K4me1 | 4.06 |
| CD4+ CD25- IL17+ PMA Ionomycin stim Th17 primary* | Immune | H3K4me3 | 4.04 |
| CD4+ CD25- Th primary* | Immune | H3K4me1 | 4.03 |
| Colonic mucosa* | GI | H3K27ac | 4.0 |
| Th1* | Immune | H3K27ac | 3.96 |
| CD4 naive primary* | Immune | H3K4me1 | 3.91 |
| CD4+ CD25int CD127+ Tmem primary* | Immune | H3K4me1 | 3.87 |
| CD8 memory primary* | Immune | H3K4me1 | 3.84 |
| Rectal mucosa* | GI | H3K4me1 | 3.74 |
| CD19 primary (UW)* | Immune | H3K4me1 | 3.72 |
| Colonic mucosa* | GI | H3K9ac | 3.63 |
| Rectal mucosa* | GI | H3K9ac | 3.57 |
| CD25- IL17- Th stim MACS* | Immune | H3K27ac | 3.54 |
| CD25int CD127+ Tmem* | Immune | H3K27ac | 3.45 |
| Th0* | Immune | H3K27ac | 3.44 |
| CD8 naive primary (UCSF-UBC)* | Immune | H3K4me1 | 3.43 |
| CD56 primary* | Immune | H3K4me1 | 3.21 |
| Rectal mucosa* | GI | H3K27ac | 3.16 |
| CD19 primary (BI)* | Immune | H3K4me1 | 2.95 |
| Treg primary* | Immune | H3K4me3 | 2.93 |
| CD8 naive primary (BI)* | Immune | H3K4me1 | 2.91 |
| CD3 primary (UW)* | Immune | H3K4me1 | 2.83 |
| CD4+ CD25- CD45RA+ naive primary* | Immune | H3K4me1 | 2.7 |
| CD3 primary (BI)* | Immune | H3K4me1 | 2.44 |
| Rectal mucosa* | GI | H3K4me3 | 2.29 |
| CD4+ CD25- IL17- PMA Ionomycin stim MACS Th primary* | Immune | H3K4me3 | 2.24 |
| Duodenum smooth muscle* | GI | H3K27ac | 2.17 |
| Duodenum Mucosa* | GI | H3K4me1 | 2.15 |
| CD34 primary* | Immune | H3K4me1 | 2.12 |

**Figure S1:**
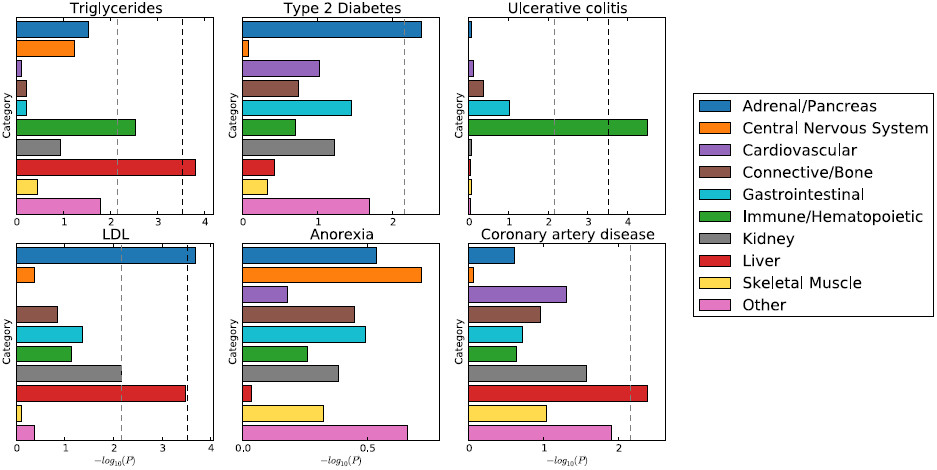
Enrichment of cell type groups for traits not included in Figure 4.The black dotted line at –log_10_(*P*) = 3:5 is the cutoff for Bonferroni significance. The grey dotted line at –log_10_(*P*) = 2:1 is the cutoff for FDR < 0.05.

